# *Tbx1* Heterozygosity in the Oligodendrocyte Lineage Shifts Myelinated Axon Composition in the Mouse Fimbria Without Behavioral Impairments

**DOI:** 10.64898/2025.12.30.697076

**Authors:** Anne Marie Wells, Takaki Tanifuji, Takeshi Takano, Arumu Endo, Gina Kang, Marisa Esparza, Qian Shi, Manzoor A. Bhat, Noboru Hiroi

## Abstract

Constitutive heterozygosity of *Tbx1*, a T-box transcription factor gene located within the 22q11.2 deletion region, results in behavioral deficits and altered composition of myelinated axons in the fimbria, together with reduced levels of an oligodendrocyte precursor cell (OPC) marker, in mice. However, the cellular origins of these effects and the extent to which axonal changes causally contribute to behavioral impairments remain unclear. We hypothesized that *Tbx1* deficiency specifically within the oligodendrocyte lineage contributes to myelin and behavioral phenotypes. To test this hypothesis, we first demonstrated through *in vitro* siRNA knockdown that *Tbx1* regulates both OPCs and mature oligodendrocytes. Subsequently, we assessed the impact of *Tbx1* heterozygosity initiated in OPCs on behavioral and myelin phenotypes in male conditional PdgfrαCre;*Tbx1*^+/flox^ mice. These mice exhibited Cre-mediated recombination in Pdgfrα-expressing brain regions and in the OPC progeny within the fimbria. At one month of age, the mutants displayed a higher rate of spontaneous alternation at the longest inter-trial interval in the T-maze compared to their wild-type littermates—an effect that was dissipated at two months. No significant phenotypic abnormalities were observed in conditional PdgfrαCre;*Tbx1*^+/flox^ mice regarding neonatal ultrasonic vocalizations, social interaction, novel object approach, anxiety-like behavior (elevated plus maze), or open-field locomotion and thigmotaxis. Electron microscopic analysis revealed a compositional shift in myelinated axons within the fimbria of adult male mutants, characterized by an increased number of myelinated axons in the 300–800 nm diameter range and a decreased number in the ∼1,200 nm and ∼1,400 nm ranges, with myelin thickness remaining unchanged across diameters. These findings indicate that *Tbx1* heterozygosity in the oligodendrocyte lineage leads to a selective shift towards smaller myelinated axons in the fimbria and a transiently higher level of capacity for working memory and cognitive flexibility. However, it does not replicate the full spectrum of myelination abnormalities or the broader cognitive and social deficits observed in constitutive *Tbx1* heterozygotes, suggesting that *Tbx1* deficiency in non-oligodendrocyte lineage cells may lead to altered myelination and neurodevelopmental behavioral impairments.

## INTRODUCTION

Cognitive and social impairments are predictors of mental illness(1, 2). Infants later diagnosed with autism spectrum disorder (ASD) exhibit delays and deviations in various aspects of motor, social, and language development(3-11). Similarly, children who subsequently develop full-scale symptoms of schizophrenia display delayed development of social cognition, working memory, attention, and processing speed(12-16), with these deficits remaining integral to the disorder following its onset(17, 18).

These dimensional deficits in mental illness have genetic underpinnings. Copy number variations (CNVs)--which consist of deletions or duplications of up to several million base pairs at specific chromosomal loci encompassing numerous protein-coding genes—are associated with high odds ratios and penetrance rates for cognitive and social deficits(19, 20) as well as mental illnesses(21-23).

Individuals with hemizygous deletions of human chromosome 22q11.2, ranging from 1.5 Mb to 3.0 Mb, exhibit cognitive, social, and emotional impairments from childhood (24-26) and are diagnosed at elevated rates with anxiety disorders, attention-deficit hyperactivity disorder (ADHD), ASD, intellectual disability (ID), and schizophrenia(27, 28). Duplication of 22q11.2 is linked to epilepsy, ID, ADHD, and ASD at rates higher than those observed in non-carriers(29-45). However, the mechanisms through which over 40 protein-coding genes in 22q11.2 contribute to dimensional deficits and clinical diagnoses remain poorly understood in humans.

Rare inherited cases of variants in the human *TBX1* gene, a T-box transcription factor located within the 22q11.2 CNV, are associated with ASD, schizophrenia, and intellectual disability in the absence of 22q11.2 CNVs(46-51). However, due to their rarity, the statistical reliability of the association between *TBX1* variants and mental illness is limited. Moreover, these patients often carry variants in other single genes throughout the genome(49), complicating the establishment of causality from these associations.

The deletion or overexpression of small murine chromosomal regions orthologous to human chromosomes or single genes provides a complementary approach (1, 52-56). Constitutive overexpression of several hundred kilobase pair segments of the mouse ortholog of 22q11.2 recapitulates distinct social and cognitive deficits associated with 22q11.2 duplication(52, 57). Larger hemizygous deletions that encompass one of these segments result in deficits in prepulse inhibition (50). These studies in mouse models suggest the presence of driver genes for specific behavioral phenotypes within these segments.

The *Tbx1* gene is located within one of these small segments. The heterozygous deletion of *Tbx1* alone results in impairments in neonatal social communication(58-60), as well as post-pubertal and adult deficits in social interaction(58), social incentive learning(61), acoustic, but not non-acoustic, prepulse inhibition(50, 61), spatial memory acquisition in the Morris water maze, and the speed to complete simple discrimination and reversal phases of attentional set-shifting, without non-specific motor speed (62). The fimbria of post-pubertal *Tbx1* heterozygous mice displays a lack of large myelinated axons, enhanced myelination of medium-sized axons, a compositional shift of myelinated axons toward smaller sizes, and a gene expression profile indicative of defective oligodendrocyte precursor cells(62). Furthermore, the volume of the secondary motor cortex and amygdala, regions involved in the vocal network(63), is reduced in post-pubertal *Tbx1* heterozygous mice (61). Additionally, virally induced overexpression of *Tbx1* in the hippocampus negatively impacts the developmental maturation of working memory capacity(64).

The cellular and developmental origins of the effects of *Tbx1* deficiency on behavioral and myelin phenotypes remain poorly understood. However, cells that express *Tbx1* are candidate substrates. Recent single-cell transcriptomic analyses have revealed that *Tbx1* is expressed in a more diverse array of cell clusters with signatures indicative of distinct cell types in the mouse than previously indicated by *in situ* hybridization and immunocytochemical studies. *Tbx1* is detectable in the neural tube, as well as endothelial cells, radial glial cells, neural progenitor cells, oligodendrocyte progenitor cells, oligodendrocytes, and various types of neurons and their progenitors (65, 66). During the embryonic period, oligodendrocyte precursor cells emerge from embryonic neural progenitor cells in the medial ganglionic eminence of the ventricular zones around E12.5, followed by a second wave from the lateral ganglionic eminence at E15.5. Oligodendrocyte precursor cells continue to be generated from neural progenitor cells in the subventricular zone during the neonatal period (67-71). Given that single-cell transcriptomic analyses can detect rare or transitional cell types, these studies suggest that *Tbx1* is present in radial glial cells and neural progenitor cells, as well as their progeny, including oligodendrocyte and neuronal lineages.

Our previous work demonstrated that *Tbx1* heterozygosity initiated in neural progenitor cells during the neonatal period recapitulates some behavioral deficits associated with constitutive *Tbx1* heterozygosity(72). As neural progenitor cells during the neonatal period give rise to oligodendrocyte precursor cells, oligodendrocytes, neural progenitor cells, and neurons(67-71), we sought to determine whether the effects of *Tbx1* heterozygosity within the oligodendrocyte lineage induces behavioral and anatomical phenotypes. Our findings indicate that *Tbx1* heterozygosity in the oligodendrocyte lineage induces a compositional shift in myelinated axons in the fimbria, while not altering myelin integrity or large myelinated axons, or impairing behavior.

## METHODS

### Mice

All protocols for animal handling and use were approved by the Institutional Animal Care and Use Committee (IACUC) of the University of Texas Health Science Center at San Antonio (UTHSCSA) in accordance with National Institutes of Health (NIH) guidelines. Mice were housed in a vivarium with a standard light phase from 7 am to 9 pm).

#### C57BL/6J mice

We used 1- and 2-month-old male C57BL6/J mice (Jax #000664, Jackson Laboratory, Bar Harbor, ME) as stimulus mice for social interaction.

#### *Pdgfrα*Cre*;Tbx1*^+/flox^ *m*ice

To conditionally induce *Tbx1* heterozygosity in oligodendrocyte precursor cells, we chose a *Pdgfrα*Cre line that initiates Cre synthesis earlier than a *Cspg4*Cre line to better induce recombination in the oligodendrocyte lineage(73). The *Pdgfrα-* promoter drives Cre-based recombination most robustly in oligodendrocyte precursor cells (74), but also in astrocytes, neurons, fibroblast-like cells, endothelial cells, vascular and leptomeningeal cells, and pericytes in the mouse brain(74-77).

We crossed hemizygous *Pdgfrα*Cre mice with *Tbx1*^+/flox^ mice to generate *PdgfrαCre;Tbx1*^+/flox^ mice. *PdgfrαCre* breeder mice (C57BL/6-Tg(*Pdgfrα*-Cre)1Clc/J; Jax #013148, Jackson Laboratory, Bar Harbor, ME) have a mixed C57BL/6J;C57BL/6N background. We backcrossed non-congenic *Tbx1*^+/flox^ mice(78) to C57BL/6J for more than 10 generations and confirmed that there was Cre-dependent recombination in a congenic *Tbx1*^+/flox^ mice(72). The genetic background of *Pdgfrα*Cre and *Tbx1*^+/flox^ lines were mixed C57BL/6J;C57BL/6N and C57BL/6J, respectively. These two C57BL/6 substrains have many behavioral differences(79). To avoid the confounding effects of their unequal genetic backgrounds(80), we used only an F1 generation derived from crosses of the *Pdgfrα*Cre and *Tbx1*^+/flox^ lines. While both male and female mice were born with all genotypes, only half as many females as males survived to complete analyses up to 2 months of age. Because the sample sizes of females tested were not sufficient for statistical reliability, our analyses were based on male mice only. No sex differences are observed in social or cognitive dimensions among carriers of 22q11.2 deletion in humans (24).

#### PdgfrαCre;ROSA-tdTomato mice

We crossed hemizygous *Pdgfrα*Cre mice (C57BL/6-Tg(*Pdgfrα*Cre)1Clc/J; Jax #013148, Jackson Laboratory, Bar Harbor, ME) with homozygous congenic *ROSA-CAG-tdTomato* mice (B6.Cg-*Gt(ROSA)26Sor^tm14(CAG-tdTomato)Hze^*/J, Jax # 007914, Jackson Lab, Bar Harbor, ME) to generate *Pdgfrα*Cre*;ROSA-CAG-tdTomato* mice. This ROSA line expresses tdTomato in the hippocampus without the action of Cre ( http://connectivity.brain-map.org/transgenic/experiment/81560256)

We determined the genotypes of the mice using our previously published protocol with primers shown in **Table S1**. The sex of the mice was determined by inspection of their external genitalia.

### *In vitro* cell culture of oligodendrocytes

Lateral ventricle tissues, including the subventricular zone (SVZ), were obtained from postnatal day 1-2 (P1-2) C57BL/6J pups. Cells were isolated in culture, with each culture derived from a single mouse. The cells were cultured in a medium (DMEM/F12, HEPES [11320–032, Gibco, Grand Island, NY, USA]) supplemented with N2 (17502048, Gibco, Grand Island, NY, USA), B27 (17504044, Gibco, Grand Island, NY, USA), epidermal growth factor (EGF) at a concentration of 20 ng/ml (AF100-15, Peprotech, Cranbury, NJ, USA). After two to four passages, the cells were dissociated from the spheres using StemPro Accutase Cell Dissociation Reagent (A1110501, Gibco, Grand Island, NY, USA). The cells were then divided into equal portions and plated on each Poly-L-ornithine (P4957, Sigma, St. Louis, MO, USA) and Bovine Fibronectin (1030-FN, R&D Systems, Minneapolis, MN, USA)-coated 6well plate (Corning, 351146, Corning, NY, USA,). The cells were cultured for 144 hours in medium with 5% fetal bovine serum after withdrawing EGF from the medium to induce and maintain differentiation. At the time of EGF withdrawal, we applied *Tbx1* siRNA (10nM, cat#4390771, s74767, Invitrogen; Thermo Fisher Scientific, Waltham, MA, USA) and control siRNA-A (10nM, cat#4390843, Invitrogen; Thermo Fisher Scientific, Waltham, MA, USA) together with siLentFect™ Lipid Reagent for RNAi, 0.5 ml (BIO-RAD, cat# 1703360, Hercules, CA, USA).

### Quantitative reverse transcription polymerase chain reaction (qRT-PCR)

In accordance with our published procedure(62), total RNA was extracted using an RNeasy Plus Mini Kit (Cat#74134, Qiagen, Germantown, USA). Complementary DNA (cDNA) was synthesized from total RNA using SuperScript IV VILO Master Mix (Cat#11766050, Invitrogen, Carlsbad, USA). qRT-PCR reactions were performed in triplicate on a QuantStudio 6 Flex Real-Time PCR System (Cat#4485694, Applied Biosystems, Waltham, USA) using the TaqMan Fast Advanced Master Mix (Cat#4444963, Applied Biosystems, Waltham, USA). We used TaqMan® Gene Expression Assays (cat# 4331182, Thermo Fisher, Waltham, MA, USA). The Taqman probes are listed in the Supplementary Material (**Table S2**). Data were analyzed using the ΔΔCt method and normalized to the reference gene *Pgk1*; we had an identical pattern of results with another reference gene *18S*.

### Whole brain tissue clearing and index matching

We used the SHIELD and SmartBatch+ (LifeCanvas Sciences, Cambridge, MA) standard pipelines for preserving and electrophoretic clearing(81) to visualize tdTomato in the brains of *Pdgfra*Cre;ROSA-CAG-tdTomato mice. We performed transcardial perfusion of 1-month old mice with ice-cold 0.9% NaCl in dH_2_O, followed by 4% paraformaldehyde (PFA) in phosphate-buffered saline (PBS). Brains were preserved using SHIELD as follows: perfused brains were 1) removed from the cranium and fixed in 4% PFA in PBS overnight at 4°C with gentle shaking; 2) incubated in SHIELD OFF solution (dH_2_O + SHIELD Buffer [cat#SH-Bf] + SHIELD Epoxy Buffer [cat#SH-Ex]) solution at 4°C with gentle shaking for 3 days; 3) incubated in pre-warmed SHIELD ON Buffer (cat#SH-ON) at 37°C for 1 day with gentle shaking; 4) incubated in the delipidation buffer (cat#DB) for 3 days at 45°C with gentle shaking. Final clearing by electrophoresis was performed with the SmartBatch+ device, with the delipidation buffer inside the clearing cup containing the samples and conduction buffer (cat#CB) for 30 h. Cleared brains were index-matched using a solution of 50% EasyIndex (RI = 1.52, cat#EI-500-1.52) + 50% dH_2_O with shaking at 37°C for 1 day, then 100% EasyIndex for one additional day until brains were transparent(81).

### Light sheet microscopy

We used a Zeiss Lightsheet 7 Microscope housed in the UTHSCSA Optical Imaging Core to perform volumetric light sheet imaging of cleared *Pdgfrα*Cre;ROSA-CAG-tdTomato brains using a pair of 5×/0.1 focusing illumination objectives and a Fluar 2.5×/0.12 detection objective at identical settings. Image tiles were stitched together using Zeiss Zen Blue software (ver 3.4). 3D volume analysis was performed with Imaris (Oxford Instruments, ver 10.02). Peak intensity of the tdTomato signal was segmented, and volumetric measurements (mm^3^) and peak intensities of the total volume (arbitrary units) were calculated for each volume in each mouse.

### Immunofluorescence

*Pdgfrα*Cre;ROSA-*tdTomato* mice were sacrificed at 1 month of age. Mice were deeply anesthetized using 4–5% isoflurane and perfused transcardially with 0.9% saline followed by 4% PFA. Brains were extracted and post-fixed in 4% PFA overnight at 4°C, followed by cryoprotection in glycerol overnight at 4°C. Coronal 60-μm thick sections were cut with a freezing microtome. For immunofluorescence labeling, sections were washed in 0.1 M PBS, blocked with 5% normal donkey serum in PBS, and incubated overnight at room temperature, as we described previously(62, 72). We used a well-validated primary rabbit antibody against myelin basic protein (1:200, MBP, ab40390, Abcam) and Donkey anti-rabbit IgG-Alexa Fluor 488 (1:200, A21206, Invitrogen), together with a nuclear marker (DAPI,1:25,000, D3571, Invitrogen). MBP is a marker of mature oligodendrocytes(82, 83). We captured images of sections that were mounted and cover-slipped with Vectashield® anti-fade mounting medium (H-1000) using a Keyence microscope (BZ-X800) microscope with BZX Hardware Module and Zeiss LSM 710 confocal microscope at the UT Health Optical Imaging Core.

#### Double staining of tdTomato with markers of oligodendrocytes

We captured a series of coronal brain sections from *Pdgfrα*Cre*;ROSA-tdTomato* mice using a Keyence BZ-X800 microscope with BZX Hardware Module and a Zeiss LSM 710 confocal microscope at the UT Health Optical Imaging Core. We used the DAPI signal to identify the anatomical boundaries of landmark structures in each image, cross-matched with equivalent levels in the Allen Brain Atlas (Allen Reference Atlas – Mouse Brain [brain atlas]. Available from atlas.brain-map.org). We then used a color scale to represent the level of tdTomato intensity within the delineated anatomical boundaries.

### Behavioral Analysis

Male neonatal mice were tested for vocalization resulting from maternal separation at P8 and P12, followed by additional behavioral tests at 1 and 2 months of age, corresponding to peripubertal and post-pubertal, adolescent periods(84), respectively; early signs of puberty begin around 1 month of age(85). Because body weight can alter behavior in mice(86) and *Pdgfrα* regulates progenitor differentiation into adipocytes(87, 88), we measured body weight at the beginning of behavioral tests. Our behavioral battery included reciprocal social interaction, novel object approach, spontaneous alternation in a T-maze, elevated plus maze, and locomotor activity and thigmotaxis in an inescapable open field(52, 57-59, 64, 89-91).

The order of the behavioral assays was based on the stress level; behaviors that occur in a home cage-like setting was given first (i.e., social interaction and novel object approach). T-maze and elevated plus maze tests permit choices and are thus considered less stressful than an inescapable open field. At least a one-day interval was provided between behavioral tests to reduce the likelihood of carryover effects(61, 62, 92), except for the T-maze, in which three different delays were given on three consecutive days.

Mice were assigned randomly to experimental groups and tested during the light phase. Experimenters were blinded to genotypes. Social interaction, novel object approach, and behaviors in the elevated plus maze were recorded using a Basler GigE camera, with video images stored in Ethovision v18 software (Noldus Information Technology, Leesburg, VA). Video images of social interaction and novel object approach were manually scored according to our established criteria. Various parameters of the elevated plus maze were automatically analyzed by Ethovision. The time point of each arm entry was recorded for spontaneous alternation, and the data were subsequently tallied manually. Locomotor activity and thigmotaxis in an inescapable open field were analyzed using Med Associates Activity Monitor 7 Software (Fairfax, VT).

### Ultrasonic vocalization

Pups were tested for vocalization induced by maternal separation at postnatal days 8 and 12. A cage containing the mother and a litter was transferred to the test room 30 min before testing. Ultrasonic vocalization was recorded for 5 min by UltraSoundGate (Avisoft, Germany) connected to a computer equipped with Avisoft-RECORDER software (Avisoft, Germany) in a test chamber (18 cm long × 18 cm wide × 30 cm high). The sampling rate and lower cutoff frequency were set at 250 kHz (format, 16-bit) and 10 kHz, respectively. A frequency window from 15–150 kHz was used for analysis. Call detection was provided by an automatic threshold-based algorithm and a hold-time mechanism (hold time = 10 ms). Sonograms were inspected, and mechanical noises were eliminated from the analysis. We used the VocalMat software, which has the lowest false positive and false negative rates(93), to determine call types. This software typically detects only the most salient components of the harmonic call type and classifies this call type differently. As the harmonic call type is often affected in genetic mouse models of neuropsychiatric disorders(58, 59, 86, 94), we manually inspected all call types and re-classified such cases as harmonic calls.

### Reciprocal Social Interaction Test

A test mouse and an age-matched unfamiliar C57BL/6J mouse as a stimulus subject were simultaneously placed in a cage (28.5 cm long × 17.5 cm wide × 12.5 cm high). Mouse behavior was recorded during two 5-min sessions with a 30-min interval between sessions. Video images were scored manually for affiliative and aggressive social interaction by a rater blinded to genotype. The following reciprocal social behaviors were scored by the duration of interaction (minimum of 1 s): aggressive (tail rattle, bite/kicks, sideway offense, boxing/wrestling), affiliative, non-aggressive (mount, pursuit, olfactory investigation, allogrooming, escape/leap), or passive (side-by-side, submissive). We analyzed the sum of time engaged in affiliative, active, and passive social interactions, but passive behavior was rarely seen in our experimental set-up (72, 86, 91, 94).

### Novel Object Approach

A test mouse was placed in a cage (28.5 cm long × 17.5 cm wide × 12.5 cm high) with an empty, non-secured 50-mL Falcon tube (3 cm diameter × 8.5 cm long) lying on its side. Videos of mouse interactions with the tube were used to manually score approach behavior by two independent analysts blinded to genotype with an inter-rater reliability of >99%. Approach behavior was scored for olfactory or non-olfactory investigations with a minimum duration of 1 s. Time spent near the tube was also analyzed.

### Spontaneous Alternation in a T-Maze

A test mouse was placed in the start site of the long arm of a black plexiglass T-maze (21.5 cm long × 10.5 cm wide × 20.5 cm high) and allowed to enter the maze and explore either the left or right short arms (31 cm long × 10.25 cm wide × 20.5 cm high) of the cage. We imposed 0 s, 15 s, or 30 s interval delays to maze reentry for 10 trials, and the three interval trials were given on three consecutive days. We manually scored the percentages of alternation and latency to reach the alternated and non-alternated arms.

### Elevated Plus Maze Test

Mice were placed in the center stage (5 × 5 cm) of the maze apparatus with four arms (30 × 5 cm) and permitted to explore two closed arms and two open arms extending from the center platform of the maze for 5 min. The maze was positioned 53 cm above the floor. The time spent in visits and the frequency of visits to the two open and two closed arms were analyzed.

### Open Field

Mice were placed in an open-field cage (27.3 cm × 27.3 cm × 20.2 cm; Med Associates, Fairfax, VT) and permitted to explore the cage for 30 min. We measured the distance and velocity of motor activity and duration of time spent in the center (19.05 cm × 19.05 cm central square) versus the margin of an open field, using the activity monitor software (Med Associates, Fairfax, VT).

### Electron Microscopy (EM)

We prepared tissue for EM to characterize myelination in the fimbria of *Pdgfrα*Cre;*Tbx1*^+/flox^ mice as we described previously(62). After behavioral assays were completed at 1 and 2 months of age, 4–5-month-old male *Pdgfrα*Cre;*Tbx1*^+/flox^ (N=3) and five control mice, including *Pdgfrα*Cre;*Tbx1*^+/+^ (N=2), wild-type (WT);*Tbx1*^+/flox^ (N=2), and WT;*Tbx1*^+/+^ (N=1), were anesthetized with 4.5% isoflurane in a chamber, and anesthesia was maintained by a nose cone with 2.0% isoflurane. The animals were transcardially perfused with 120 mL of 0.9% saline followed by fixation with 120 mL of 0.1 M buffer (pH 7.4; cat#11653 Electron Microscopy Science, Hatfield, PA) with 2.5% sodium glutaraldehyde (cat#16310, Electron Microscopy Sciences, Hatfield, PA) and 2.5% paraformaldehyde (cat#19202, Electron Microscopy Sciences, Hatfield, PA). Brains were extracted and post-fixed in the same solution at 4°C for 7 weeks. The fimbria from the two hemispheres were obtained separately using a vibratome and placed in 0.1 M sodium cacodylate buffer overnight. Tissues were rinsed with 0.1 M sodium cacodylate buffer to remove aldehydes and placed in 2% osmium tetroxide solution (OsO_4_; cat#19150, Electron Microscopy Sciences, Hatfield, PA) in 0.1 M sodium cacodylate buffer for 1 h. Tissues were dehydrated in a series of ethanol solutions and embedded in molds containing Polybed resin (Poly/Red® 812 Embedding media, cat#08791-500 Polysciences, Inc., Warrington, PA). We cut 100-nm sections and collected them on square 150-mesh copper grids (cat#7551C, Polysciences, Inc., Warrington, PA) and stained them with uranyl acetate (7 g in 100 mL deionized water)/Reynold’s lead citrate (1.33 g lead nitrate,1.76 g sodium citrate in 30 mL triple-distilled water, 8 mL 1 N NaOH). Uranyl acetate stains membranous structures and structures containing nucleic acid; lead citrate binds to RNA-containing structures and the hydroxyl groups of carbohydrates.

Each fimbria section was viewed in the EM grid squares. Images were screened at 1,000× magnification. For each section, we used all grid images that contained fimbria tissue with round axons without wrinkles, folds, or tears in the sections. One image was captured at the center of each grid field at 20,000× magnification. Images were analyzed by MyelTracer(95). Only axons whose myelin was fully within the image were analyzed quantitatively.

We obtained multiple images for each animal from a section of the right and left hemispheres. The positions of sections sliced from the fimbria varied between the two hemispheres and did not necessarily match the two sides; in one case, only one hemisphere was available because of the accidental loss of a tissue slice. The number of images varied across tissue slices, resulting in unequal image numbers. Many axons from each grid square and many grid squares from each hemisphere were analyzed for each animal. As the number of images and positions of each image varied between the two hemispheres and did not match between the two sides, the hemisphere could not be used as a replicate. Thus, we pooled data for each animal as a technical replicate; data points within and across images were technical replicates. As there was considerable variability in the data, we could not average the values for each animal. The averages of the total axon diameter per genotype were also not appropriate for this data set, as axon numbers differed between genotypes at specific axon diameters (≥300 nm to <400 nm; ≥1,200 nm to <1300 nm) (genotype × axon diameter range, F(27,5880) = 6.336, *P* <1.0 × 10^−4^). Thus, linear mixed models were used, as reported previously(62). Data were assessed for the number of myelinated axons, the thickness of myelination, and the g-ratio along a 100 nm unit axon diameter. Data from different images and hemispheres were embedded in random models.

### Statistical Analysis

We used SPSS (v29.0.2.0 (20), IBM Corporation) to perform all statistical analyses. Among-group and between-group comparisons of the data were performed using analysis of variance (ANOVA) and Student’s two-tailed t-test (α = 0.05). We determined normality and homogeneity of variance using the Shapiro-Wilk test and Levene’s homogeneity of variance test, respectively If either assumption was violated, we analyzed the data using a linear mixed model, Kruskal Wallis tests, or Mann–Whitney U tests. The Greenhouse–Geisser correction was applied if sphericity was violated and the estimated epsilon was less than 0.75. If multiple tests were applied to a data set, the significance level was adjusted using the Benjamini–Hochberg correction, with a false discovery rate of 5%. We used GraphPad Prism software (v9; GraphPad Software, San Diego, CA) to generate all graphs. All statistical analyses are presented in Table S3.

## Data Availability

All data that support the findings and conclusions are provided within the article. All raw data and additional information are available upon request.

## List of abbreviations

CNVs: Copy number variations
ASD: Autism Spectrum disorder
scRNA-seq: single cell RNA-seq
Pdgfra: Platelet-derived growth factor receptor alpha
MAG: Myelin associated glycoprotein
MOG: Myelin oligodendrocyte glycoprotein,
MBP: myelin basic protein

## Declarations

### Ethics approval and consent to participate

All protocols for animal handling and use were approved by the Institutional Animal Care and Use Committee (IACUC) of the University of Texas Health Science Center at San Antonio (UTHSCSA) in accordance with National Institutes of Health (NIH) guidelines.

### Consent for publication

Not applicable

### Availability of data and materials

All data and materials are available upon request.

### Competing interests

The authors declare no competing interests.

### Funding

T. Takano, GK, ME, TH, and NH were supported by the National Institute of Health (R01MH099660; R01DC015776). T. Tanifuji was supported by SENSHIN Medical Research Foundation and Uehara Memorial Foundation. QS and MAB were supported by the National Institute of Health (R01GM063074). AMW was funded by F30MH134482, UT Health San Antonio CTSA T32 (T32R004545), UT Health San Antonio Neuroscience T32 (T32NS082145), STX-MSTP (NIH T32GM113896/T32GM145432), and SfN NSP Fellowship (R25NS089462). The Zeiss Lightsheet 7 microscope was funded by the NIH S10 grant 1S10OD030383. The content is solely the responsibility of the authors and does not necessarily represent the official views of the National Institutes of Health.

### Authors’ contributions

Anne Marie Wells: Conducted lightsheet brain analysis, perfused mice and applied MyelTracer to images for electron microscopy, genotyping, social interaction assessments, and manuscript writing.

Takaki Tanifuji: Performed qRT-PCR analysis, immunofluorescent staining, confocal imaging and analyses, electron microscopy image capturing, and figure preparation.

Takeshi Takano: Conducted statistical analyses and prepared all figures except for immunofluorescent images.

Arumu Endo: Performed qRT-PCR analysis and figure preparation.

Gina Kang: Conducted all behavioral testing.

Marisa Esparza: Managed breeder maintenance, ensured quality control of data input, performed statistical analyses, conducted mouse perfusion, and carried out genotyping.

Qian Shi: Responsible for electron microscopy preparation and manuscript writing.

Manzoor A. Bhat: Engaged in electron microscopy preparation and manuscript writing.

Noboru Hiroi: Designed all experiments, supervised all personnel, prepared electron microscopy samples, captured electron microscopy images, performed immunofluorescent staining and prepared figures, and wrote the manuscript.

## Acknowledgments

The LifeCanvas device was purchased with generous donations by 10 scientists at UT Health San Antonio. We thank Dr. Bernice Morrow for providing *Tbx1*^+/flox^ breeders, Ms. Monica D. Alarcon for technical assistance with EM analysis, Dr. Lacey B. Sell for her help with MyelTracer software, and Dr. Takeshi Hiramoto for valuable comments.

## RESULTS

### Effects of *Tbx1* Knockdown on Markers of Oligodendrocyte Lineage *In Vitro*

We aimed to identify the earliest stage in the oligodendrocyte lineage affected by *Tbx1* deficiency. To achieve this aim, we developed an *in vitro* screening assay utilizing neonatal neural progenitor cells derived from the subventricular zone of C57BL/6J pups sacrificed on postnatal day 2, which are capable of generating both neuronal precursor cells and oligodendrocyte precursor cells(71). We assessed the expression of *Cspg4*, a marker for oligodendrocyte precursor cells, as well as *Mag, Mbp, Mog*, and *Plp1*, which are indicative of maturing and mature myelinating oligodendrocytes (67), at 144 hours after EGF was withdrawn (i.e., differentiation) and *Tbx1* siRNA was applied.

The application of *Tbx1* siRNA resulted in a significant reduction in the mRNA levels of *Tbx1, Cspg4*, and all markers associated with mature oligodendrocytes, with *Pgk1* (**Figure 1**) and *18S* (**Table S3-Figure 1**) as reference genes. These *in vitro* findings align with our *in vivo* data, which demonstrate that *Cspg4* (also referred to as *Ng2*) was diminished in the fimbria of *Tbx1* heterozygous mice(62).

**Figure 1.**
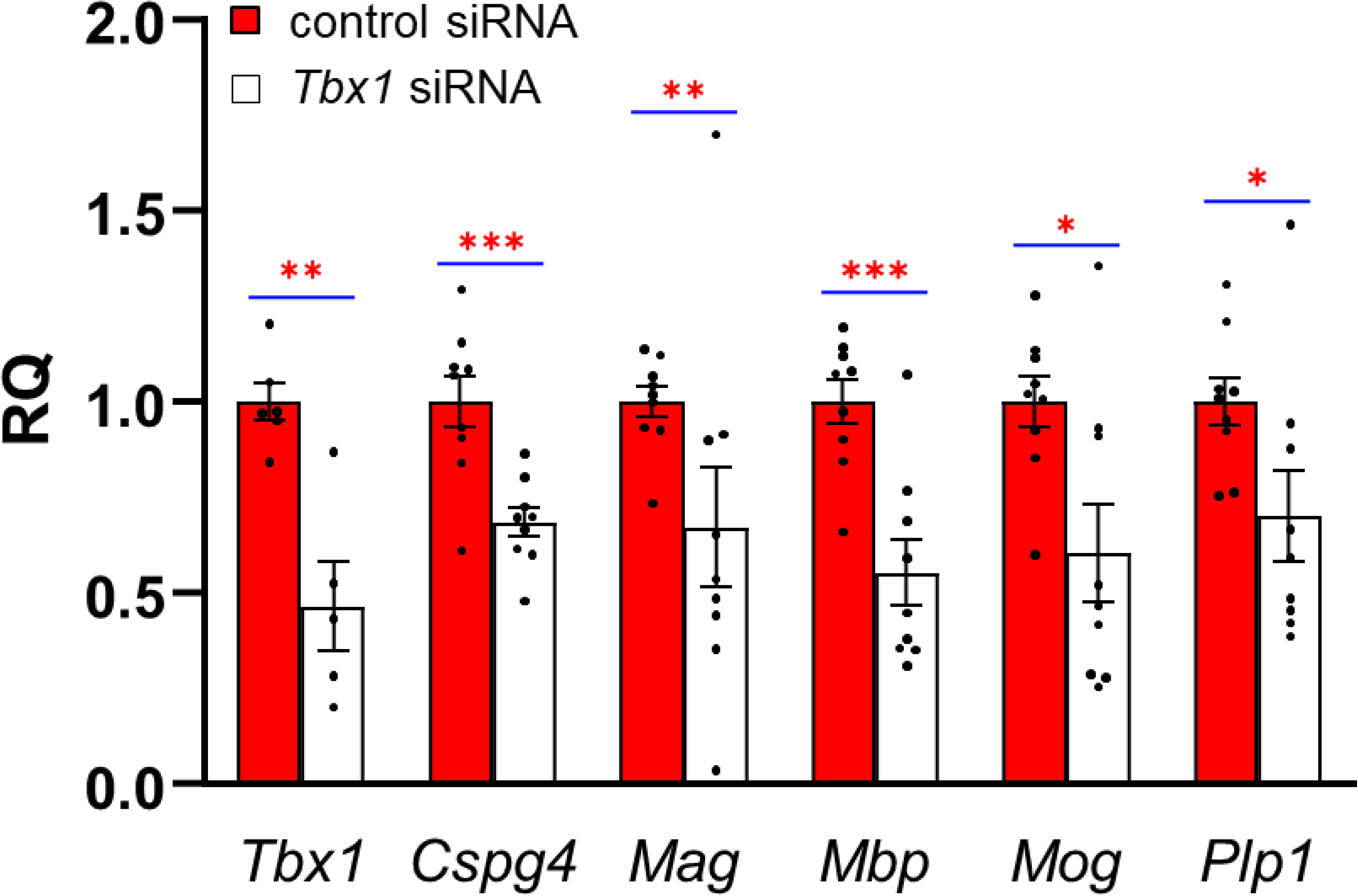
The relative quantification (RQ) (mean ± SEM) of expression levels at each stage of oligodendrogenesis, as determined by quantitative reverse transcription polymerase chain reaction (qRT-PCR) using *Pgk1* as a reference gene. Three cell clones were derived from three different C57BL/6J pups at postnatal days 2 or 3, with data obtained from three culture wells for each clone, except for one *Tbx1* siRNA-treated clone where *Tbx1* mRNA was undetectable and thus excluded from the analysis. A linear mixed model showed that Tbx1 siRNA reduced expression of all genes tested equally (Treatment, F(1,89)=14.665, p=0.0002386; Gene, F(5,89)=0.8440, p=0.5222; Treatment x Gene, F(5,89)=0.8440, p=0.5222). Mann-Whitney tests confirmed that *Tbx1* siRNA reduced expression of each gene. Statistically significant differences between control siRNA and *Tbx1* siRNA groups are indicated by *, **, and *** for p <0.05, p<0.01, and p<0.005, respectively; they remained significant following Benjamini-Hochberg correction at a 5% false discovery rate (FDR).

Given that our culture includes both neuronal (58, 72) and oligodendrocyte (see **Figure 1**) lineages, as well as residual neural progenitor cells, the mechanism by which how *Tbx1* knockdown alters the expression of myelin marker genes remains unclear. It is possible that *Tbx1* knockdown in oligodendrocyte precursor cells leads to a cell-autonomous reduction in myelin markers; alternatively, reduced *Tbx1* levels in neonatal neural progenitor cells or their neuronal progeny may indirectly influence the expression of these oligodendrocyte markers.

### Localization of Recombination in *Pdgfrα-Cre*;ROSA-tdTomato Mice

Our data indicate that *Tbx1* deficiency begins to impact the stage of oligodendrocyte precursor cells (see **Figure 1**). Consequently, we planned to initiate *Tbx1* heterozygosity in oligodendrocyte precursor cells using a conditional heterozygous mouse model. This mouse employed Cre-based initiation of *Tbx1* heterozygosity, guided by the promoter of *Pdgfrα*, a gene expressed in oligodendrocyte precursor cells of both embryonic and post-embryonic origins.

To identify the brain regions and cell types where *Tbx1* recombination is anticipated, we utilized PdgfrαCre;ROSA-tdTomato mice as the best available proxy. Alternative methodologies are not applicable to *Tbx1*. Recent single-cell RNA sequencing analyses reveal that *Tbx1* is present in proliferating cells, including neural progenitor cells, lung AT2 cells, spermatogonia, spermatids, and embryonic mesenchymal cells(65, 66). However, although *Tbx1* mRNA and protein products are expressed in actively proliferating neural progenitor cells, their levels significantly decrease in differentiated cells (72). As a result, the levels of *Tbx1* mRNA and protein products in our conditional *Tbx1* heterozygous mice are expected to reflect reductions due to cell-cycle changes, recombination-induced heterozygosity, or a combination of both factors.

Detecting TBX1 protein in the mouse brain presents challenges, with notable exceptions in endothelial cells(96) and neonatal neural progenitor cells(58, 72) partly due to epitope masking caused by fixation(58). Additionally, approaches to localize TBX1 using reporters under the endogenous *Tbx1* gene or its promoter have not yielded detectable signals in cells other than endothelial cells within the mouse brain(50, 96, 97), making it difficult to assess its colocalization with Pdgfrα protein.

In theory, Pdgfra-positive and MBP-positive cells could be isolated via fluorescence-activated cell sorting (FACS). However, while *Tbx1* is present in oligodendrocyte precursor cells and immature oligodendrocytes, its expression is notably low in mice(65, 66). Evaluating further reduced *Tbx1* levels is technically challenging due to the floor effect in heterozygous mice.

We first assessed the extent of *PdgfrαCre*-mediated recombination throughout the brain, employing a tissue clearing technique (SmartBatch+) and utilizing lightsheet microscopy to visualize the 3D volume of tdTomato signals in the brains of PdgfrαCre;ROSA-tdTomato mice (**Figure 2A**). The highest levels of tdTomato expression were observed in a pair of arch-shaped structures located along the lateral and third ventricles. Additionally, clusters of tdTomato signals were detected on the ventral surface of the posterior brain, while lower levels of signals were observed throughout other regions.

**Figure 2.**
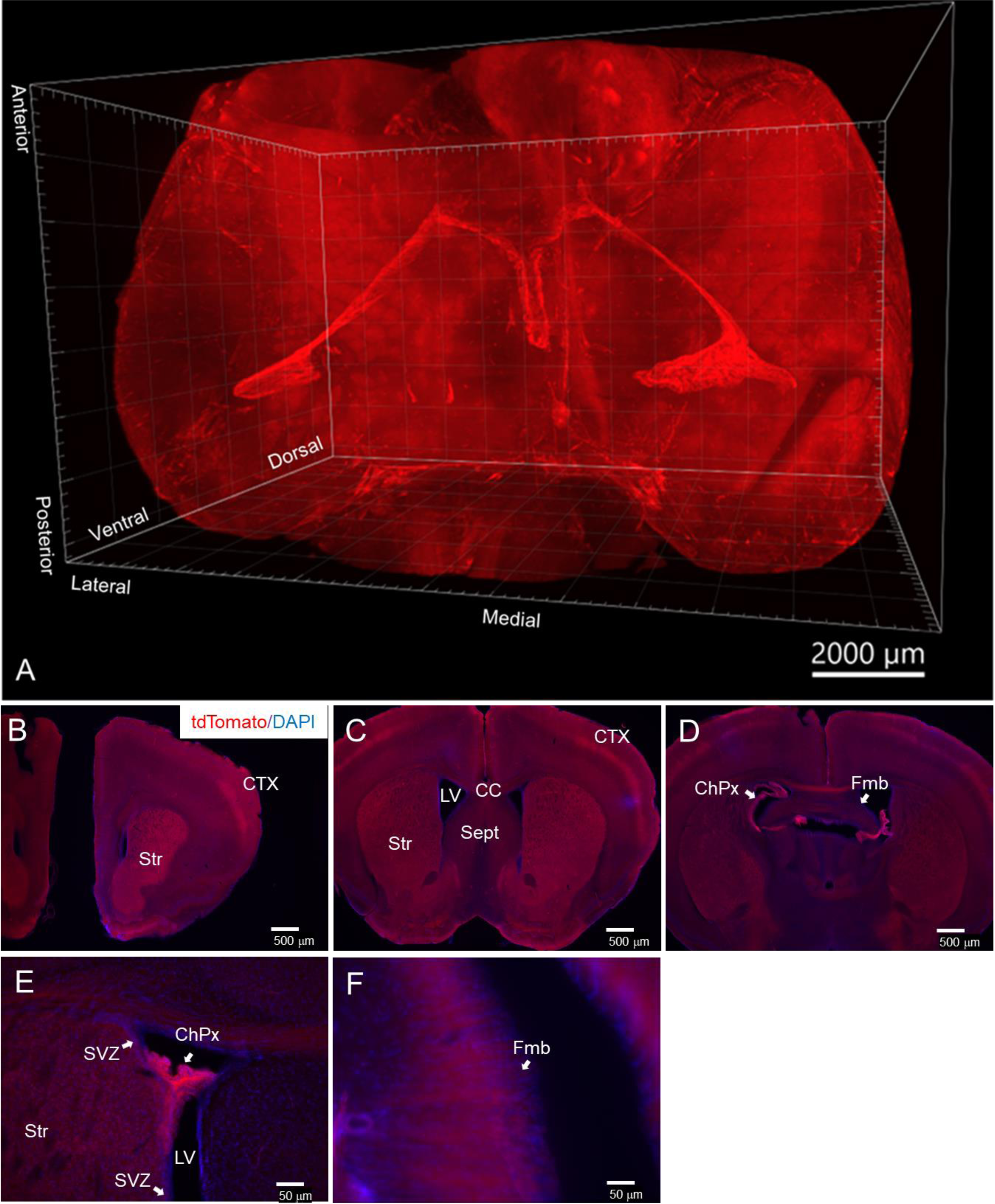
**A**. A representative lightsheet image of the brain from a 1-month-old male PdgfrαCre;ROSA-tdTomato mouse. Following SmartBatch+ tissue delipification, tdTomato signals were visualized using a Zeiss LSM 7 Lightsheet microscope with a 2.5x objective (N = 3). **B-D**. tdTomato expression in 1-month-old PdgfrαCre;ROSA-tdTomato mice is depicted together with blue DAPI signals at the levels of the anterior striatum (**B**), middle striatum (**C**), posterior striatum/fimbria (**D**), anterior lateral ventricle (**E**), middle lateral ventricle (**F**), and fimbria (**H**). Abbreviations: ChPx, choroid plexus; LV, lateral ventricle; Str, striatum; Sept, septum; CC, corpus callosum; CTX, cortex; HGCL, hippocampal granule cell layer; fmb, fimbria; SVZ, subventricular zone. N = 3.

To identify brain regions exhibiting tdTomato signals in PdgfrαCre;ROSA-tdTomato mice, we analyzed a series of coronal sections from 1-month-old PdgfrαCre;ROSA-tdTomato mice. High levels of tdTomato signals were present in the striatum, superficial layers of the neocortex (**Figure 2BC**), choroid plexus within the lateral ventricle (**Figure 2ED**) and the fimbria (**Figure 2F**). This expression pattern, driven by the *Pdgfrα* promoter, aligns with previously reported distributions of *Pdgfrα* mRNA and protein, as well as recombination activity associated with the *Pdgfrα* promoter in the mouse brain (98-100). Moreover, the intense staining of the choroid plexus is consistent with the prominently tdTomato+ structures resembling the shapes of the lateral and third ventricles (see **Figure 2A**).

Embryonic Pdgfra-positive cells give rise to distinct pre-oligodendrocyte precursor cells by the perinatal period; however, they exhibit similar profiles during the postnatal and adult periods(67, 101). PdgfraCre is expected to induce recombination in oligodendrocyte precursor cells(74). Once tdTomato is expressed through recombination in the PdgfraCre;ROSA-tdTomato mouse line, it remains expressed in their progeny, including mature oligodendrocytes and myelinated fibers. Given that myelination of the fimbria is selectively affected in constitutive *Tbx1* heterozygous mice(62), we examined whether tdTomato colocalized with MBP, a marker of mature oligodendrocytes and myelinated fibers, in the fimbria(102). MBP-positive fibers were present throughout the fimbria (**Figure 3AB**). TdTomato signals were more prominent laterally than medially (**Figure 3C**) and were colocalized with MBP-positive fibers (**Figure 3D**) in the fimbria. These findings confirm that tdTomato, activated by PdgfraCre in PdgfraCre;ROSA-tdTomato mice, is present in myelinated fibers in the fimbria.

**Figure 3.**
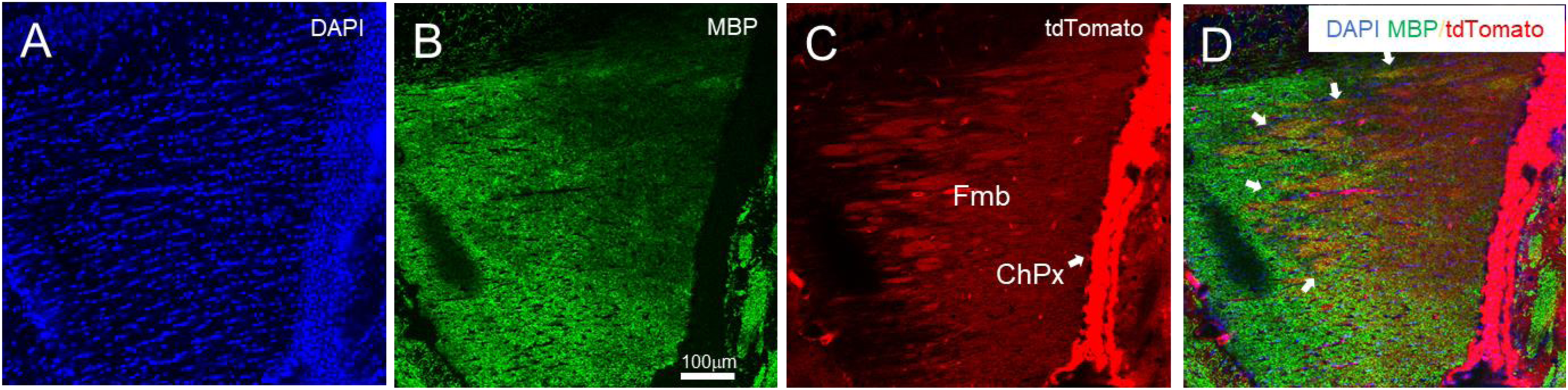
A Representative Case of Confocal Images of the Fimbria of a 1-Month-Old Male PdgfrαCre;ROSA-tdTomato Mouse. Sections were stained for DAPI (**A**, blue) and MBP (**B**, green). **C**. Red tdTomato signals originated from the PdgfrαCre;ROSA-tdTomato genotype. **D**. A composite image of the three colors is presented; white arrows indicate an area where fibers are positive for both MBP and tdTomato. N = 4. ChPx, choroid plexus; fmb, fimbria.

Intense tdTomato signals were evident in the choroid plexus located laterally to the lateral surface of the fimbria (**Figure 2D**). The cell types in the mouse choroid plexus that express *Pdgfra* include fibroblasts, mural cells, and endothelial cells during embryonic development, as well as fibroblasts and macrophages during the post-embryonic period in the mouse brain. Although gene profiles indicative of oligodendrocyte precursor-like cells are detectable during the embryonic period, their precise location within the choroid plexus remains unclear (66, 103-105). The robust tdTomato signal observed in the choroid plexus at 1 month of age likely originates from some or all of these cell types and any developmental time point. It is important to note that tdTomato signals detected at one month of age reflects recombination that has occurred any developmental time prior to this age.

### Effects of *Tbx1* Heterozygosity in Oligodendrocyte Precursor Cells on Behavior

The growth rate may influence behavior; PdgfrαCre;*Tbx1*^+/flox^, referred to as o*Tbx1*^+/−^ mice, may exhibit developmental delays, with their phenotypes potentially reflecting a lag in development compared to their wild-type littermates. However, neonatal o*Tbx1*^+/+^ and o*Tbx1*^+/−^ pups demonstrated comparable increases in body weight (**Figure S1**).

To investigate the impact of *Tbx1* heterozygosity in oligodendrocyte precursor cells and their progeny on behavior, mice were assessed tested for social communication during the neonatal period and subjected to a series of tests evaluating social, cognitive, anxiety-related, and motor behaviors at one and two months of age. This timing aligns with the peal of myelination, which occurs between the fourth and fifth postnatal weeks in rodents(106, 107) and the onset of puberty around 1 month of age and adolescence around 2 months of age(84). Constitutive *Tbx1* heterozygous mice exhibit impairments in 1) neonatal social communication as early as P7, 2) social behaviors, 3) working memory/cognitive flexibility in a T-maze and attentional set shifting, and 4) affect-related behaviors in an inescapable open field, although not in an elevated plus maze during adolescence(58-62).

We first assessed differences among the three control groups: PdgfraCre;*Tbx1*^+/+,^ WT;*Tbx1*^+/flox^, and WT;*Tbx1*^+/+^. No significant differences were observed among the controls for any behavioral measure, expect at three time points regarding total margin time in the open field (see **Table S3-Figure 8B**). When analyzing all four genotypes together, a significant genotype effect was found without an interaction effect with time (Genotype, p=0.0256; Genotype x Time, p=0.665). However, post-hoc comparisons did not reveal significant differences between WT;*Tbx1*^+/flox^ and WT;*Tbx1*^+/+^ mice only at 5 min and 15 min (see **Table S3, Figure 8B**). Given the sample size for these two control groups was n=6 each, we lacked confidence in these statistical values. Consequently, combined all the control genotypes into a single control group, designated as *oTbx1*^+/+^, for comparison with *oTbx1*^+/−^ (66).

Constitutive *Tbx1*^+/−^ mice display various neonatal, peri-adolescent, and postnatal behavioral phenotypes(58, 59). Thus, we evaluated mice for neonatal ultrasonic vocalization on P8 and P12–P13 and conducted additional tests with peripubertal (1 month) and adolescent (2 months) mice, allowing 1–2 day intervals between tasks to minimize potential carryover effects from prior testing(92).

We found no differences among o*Tbx1*^+/−^ pups regarding the number, percentage, and duration of various neonatal vocal call types on P8 and P12 (**Figure 4A–C**). At 1 and 2 months of age, mice were sequentially assessed for social interaction, novel object approach, spontaneous alternation in a T-maze, anxiety-related behavior in an elevated plus maze, and locomotor activity and thigmotaxis in an inescapable open field. *oTbx1*^+/−^ and o*Tbx1*^+/+^ mice were indistinguishable in social interaction (**Figure 5A**) and novel object approach (**Figure 5B**). In a T-maze, o*Tbx1*^+/−^ mice at 1 month of age exhibited higher spontaneous alternation rates at the longest delay (**Figure 6A**); otherwise, o*Tbx1*^+/+^ and o*Tbx1*^+/−^ mice were indistinguishable in rates of spontaneous alternation at 2 months of age and in latencies to correct choices at both ages (**Figure 6B–D**).

**Figure 4.**
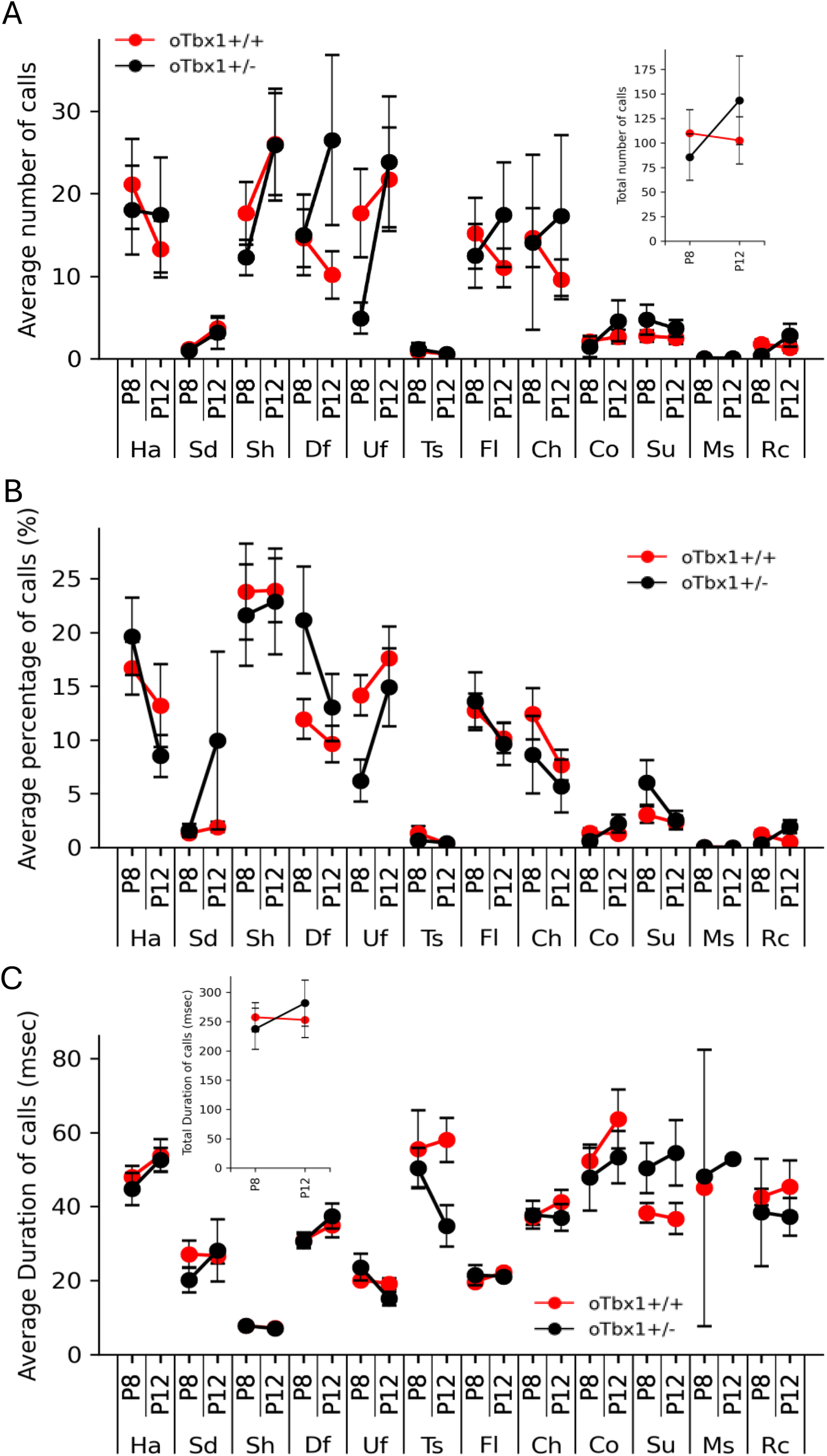
The number (**A**), proportion (**B**), and duration (**C**) (mean ± standard error of the mean [SEM]) of 12 call types emitted on postnatal days (P) P8 and P12. The two genotypes exhibited differences in certain interaction effects: **A**. Number: Genotype, F(1,912)=0.1966, p = 0.6576; Genotype x Postnatal day, F(1, 912)=4.835, p = 0.0281; Genotype x Call type, F(11, 912)=0.7077, p = 0.7319; Genotype x Postnatal day x Call types, F(11, 912)=0.3686, p = 0.9679. **B**. Proportion: Genotype, F(1,912)=0.0107, p = 0.9176; Genotype x Postnatal day, F(1, 912)=0.0107, p = 0.9176; Genotype x Call type, F(11, 912)=1.3657, p = 0.1837; Genotype x Postnatal day x Call types, F(11, 912)=0.7966, p = 0.6437. **C**. Duration: Genotype, F(1,560)=0.0435, p = 0.8349; Genotype x Postnatal day, F(1, 560)=0.10519, p = 0.7458; Genotype x Call type, F(11, 560)=3.361, p = 0.00016; Genotype x Postnatal day x Call types, F(11, 560)=1.0389, p = 0.4101. However, Mann-Whitney U tests did not reveal any significant genotype effects at any postnatal day for any call type after applying Benjamini-Hochberg corrections at 5% false discovery rate (FDR). **Inset** in **A**: The total number of ultrasonic vocalizations (mean ± SEM) emitted at P8 and P12. The two genotypes did not differ on the two postnatal days (Genotype, F(1,76)=0.1559, p = 0.694; Genotype x Postnatal day, F(1, 76)=1.103, p = 0.2969). **Inset** in **C**: Duration: Genotype, F(1,32)=0.0002, p = 0.9878; Genotype x Postnatal day, F(1, 32)=0.3617, p = 0.5518). Ha, harmonic; Sd, step-down; Sh, short; Df, down frequency modulation; Uf, up frequency modulation; Ts, two steps; Fl, flat; Ch, chevron; Co, complex; Su, step-up; Ms, multiple steps; Rc, reverse chevron. *oTbx1*^+/+^ mice, N = 27; *oTbx1*^+/−^ mice, N = 11 at P8; *oTbx1^+/+^* mice, N = 30; *oTbx1*^+/−^ mice, N = 12 at P12.

**Figure 5.**
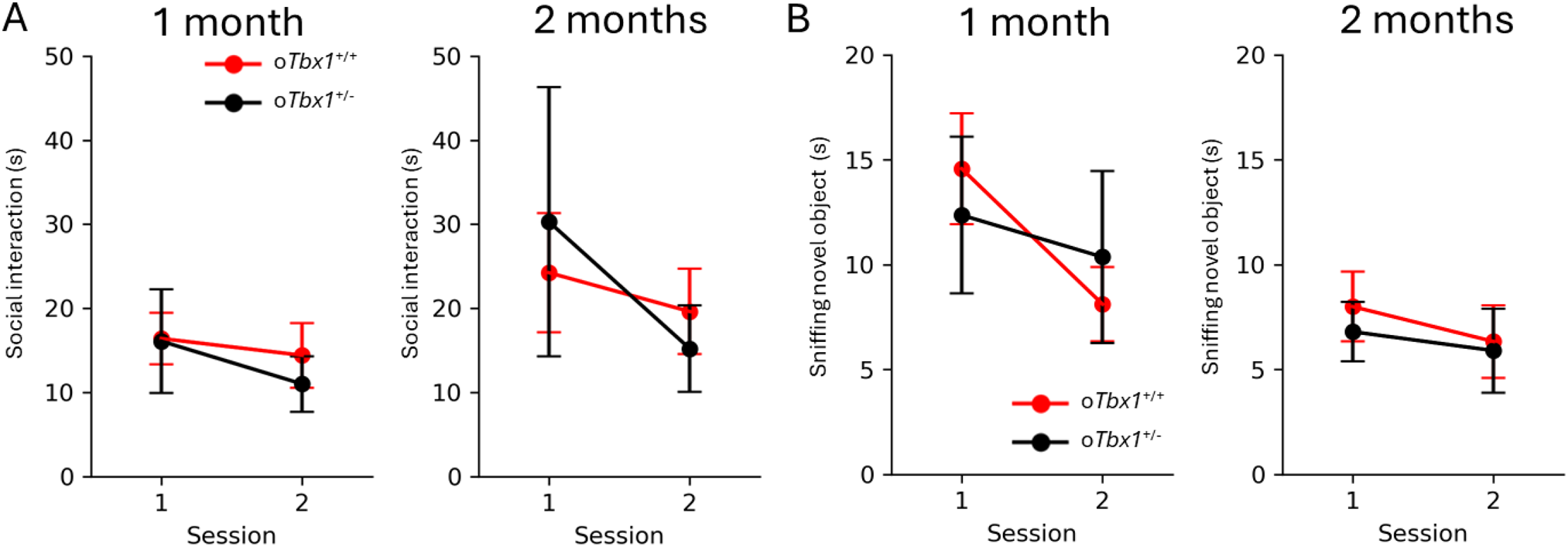
Interaction with Social and Non-Social Stimuli. **A**) Social Interaction. The *oTbx1*^+/−^ (PdgfrαCre;Tbx1^+/flox^) mice and *oTbx1*+/+ mice did not differ in the amount of time (mean ± SEM) spent in affiliative social interaction (1 month: genotype, F(1, 72) = 0.1118, p = 0.739; genotype × session, F(1, 72) = 0.262, p = 0.6102; 2 months: genotype, F(1, 58) = 0.00547, p = 0.9413; genotype x session, F(1, 58) = 1.2175, p = 0.2744). The three genotypes of *oTbx1*+/+ (1 month, N = 8, PdgfrαCre;*Tbx1*^+/+^, N = 10, Wild-type;*Tbx1*^+/+^, N = 9, Wild-type;*Tbx1*^+/flox^; 2 months, N = 8, PdgfrαCre;*Tbx1*^+/+^, N = 6, Wild-type;*Tbx1*^+/+^, N = 7, Wild-type;*Tbx1*^+/flox^) did not show significant differences (1 month: genotype, F(2, 48) = 1.686, p = 0.1960; genotype × session, F(2, 48) = 0.843, p = 0.4366; 2 months: genotype, F(2, 36) = 0.0392, p = 0.9616; genotype x session, F(2, 36) = 0.2382, p = 0.7893), and were thus combined as *oTbx1*+/+. *oTbx1*^+/+^ mice, N = 27; *oTbx1*^+/−^ mice, N = 11 at 1 month; *oTbx1*^+/+^ mice, N = 21; *oTbx1*^+/−^ mice, N = 10 at 2 months. **B**) Approach to a novel, non-social object. The two genotypes did not differ in the amount of time spent in affiliative social interaction (1 month: genotype, F(1, 70) = 2.2507E-05, p = 0.9962; genotype × session, F(1, 70) = 1.0305, p = 0.31354; 2 months: genotype, F(1, 58) = 0.11634, p = 0.7342; genotype × session, F(1, 58) = 0.0821, p = 0.77545). The three genotypes of *oTbx1*^+/+^ (1 month: N = 8, PdgfrαCre;*Tbx1*^+/+^, N = 9, Wild-type;*Tbx1*^+/+^, N = 9, Wild-type;*Tbx1*^+/flox^; 2 months: N = 8, PdgfrαCre;*Tbx1*^+/+^, N = 6, Wild-type;*Tbx1*^+/+^, N = 7, Wild-type;*Tbx1*^+/flox^) did not differ (1 month: genotype, F(2, 46) = 0.1772, p = 0.8382; genotype × session, F(2, 46) = 0.9318, p = 0.4012; 2 months: genotype, F(2, 36) = 0.0580, p = 0.9437; genotype × session, F(2, 36) = 0.6581, p = 0.5239) and were therefore combined as *oTbx1*^+/+^. *oTbx1*^+/+^ mice, N = 26; *oTbx1*^+/−^ mice, N = 11 at 1 month; *oTbx1*^+/+^ mice, N = 21; *oTbx1*^+/−^ mice, N = 10 at 2 months.

**Figure 6.**
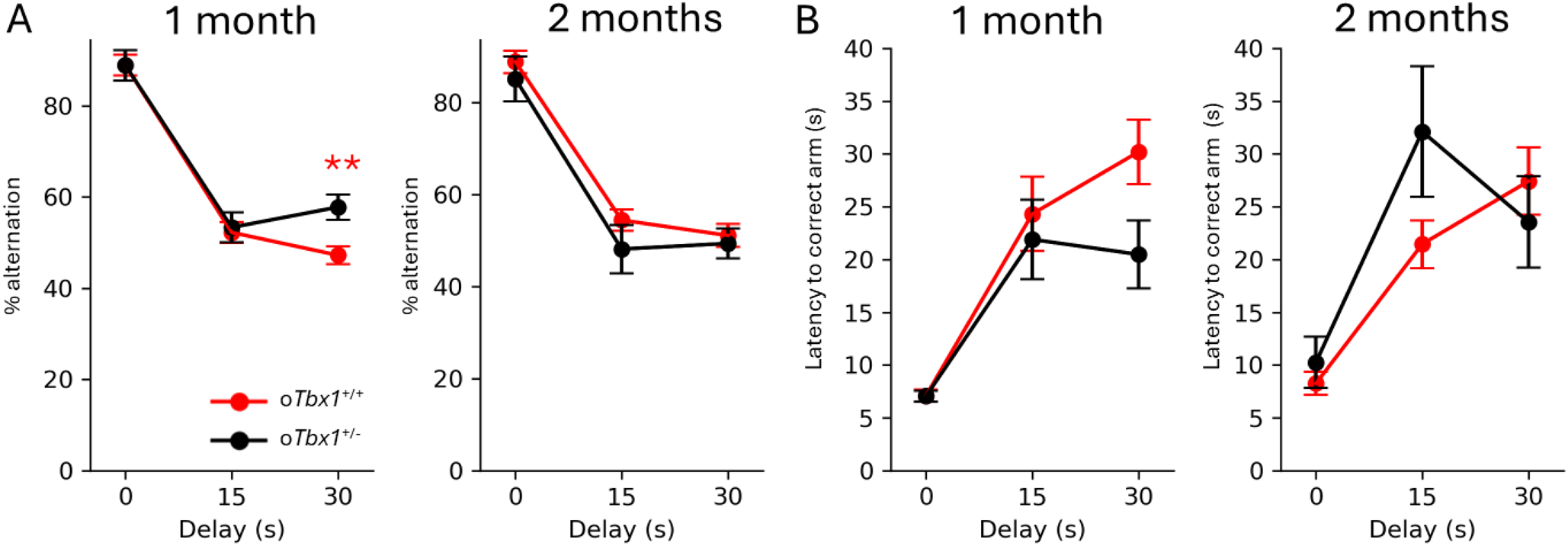
Spontaneous alternation in a T-maze. Compared to male *oTbx1*^+/+^ (N = 20) mice, male *oTbx1*^+/−^ (N = 10) mice performed at higher percentages of correct spontaneous alternations (mean ± SEM) at the longest delay (30 s) in a T-maze at 1 month of age, but not at 2 months. Specifically, at 1 month: % alternation, genotype, F(1, 96) = 4.66, p = 0.0333; genotype × delays, F(2,96) = 3.1509, p = 0.0473; at 2 months: genotype, F(1, 81) = 2.047, p = 0.1564; genotype × delays, F(2, 81) = 0.2439, p = 0.7841. The three genotypes of *oTbx1*^+/+^ (1 month: N = 8, PdgfrαCre;*Tbx1*^+/+^, N = 7, Wild-type;*Tbx1*^+/+^, N = 9, Wild-type;*Tbx1*^+/flox^; 2 months: N = 8, PdgfrαCre;*Tbx1*^+/+^, N = 6, Wild-type;*Tbx1*^+/+^, N = 6, Wild-type;*Tbx1*^+/flox^) did not differ (1 month: genotype, F(2, 63) = 0.1046, p = 0.9008; genotype × delays, F(4, 63) = 0.2648, p = 0.8995; 2 months: genotype, F(2, 51) = 0.1197, p = 0.8874; genotype × delays, F(4, 51) = 0.6401, p = 0.6363) and consequently were combined as *oTbx1*^+/+^. *oTbx1*^+/−^ mice exhibited higher rates of correct alternations at the 30-second delay compared to *oTbx1*+/+ mice, as determined by the Mann-Whitney test (p = 0.0027). **B**) The two groups did not exhibit significant differences in the latency to correct choices in the T-maze at 1 month (genotype, F(1, 96) = 2.181, p = 0.1430; genotype × delays, F(2, 96) = 2.058, p = 0.1332) or at 2 months (genotype, F(1, 81) = 1.1135, p = 0.2945; genotype × delays, F(2, 81) = 2.5825, p = 0.0818). The three genotypes of *oTbx1*^+/+^ (1 month, N=8), PdgfrαCre;*Tbx1*^+/+^ (N=7), Wild-type;*Tbx1*^+/+^ (N=9), and Wild-type;*Tbx1*^+/flox^ (2 months, N=8), PdgfrαCre;*Tbx1*^+/+^ (N=6), Wild-type;*Tbx1*^+/+^ (N=6), and Wild-type;*Tbx1*^+/flox^, demonstrated a significant interaction effect at 1 month of age (genotype, F(2, 63) = 1.5787, p = 0.2143; genotype × session, F(4, 63) = 4.5625, p = 0.00267), but no significant interaction at 2 months of age (genotype, F(2, 51) = 0.1631, p = 0.8500; genotype × session, F(4, 51) = 1.0611, p = 0.3853). However, Mann-Whitney U tests revealed that none of the three genotypes significantly differed from one another at 1 month of age after applying the Benjamini-Hochberg correction (p>0.05 for all pairs). The three genotypes were combined as o*Tbx1*^+/+^. The sample sizes were as follows: *oTbx1*^+/+^ mice, N = 24; *oTbx1*^+/−^ mice, N = 10 at 1 month; *oTbx1*^+/+^ mice, N = 20; *oTbx1*^+/−^ mice, N = 9 at 2 months.

The two genotypes were indistinguishable in the percentage of time spent in open arms (**Figure 7A**) and visits to open arms (**Figure 7B**) of the elevated plus maze.

**Figure 7.**
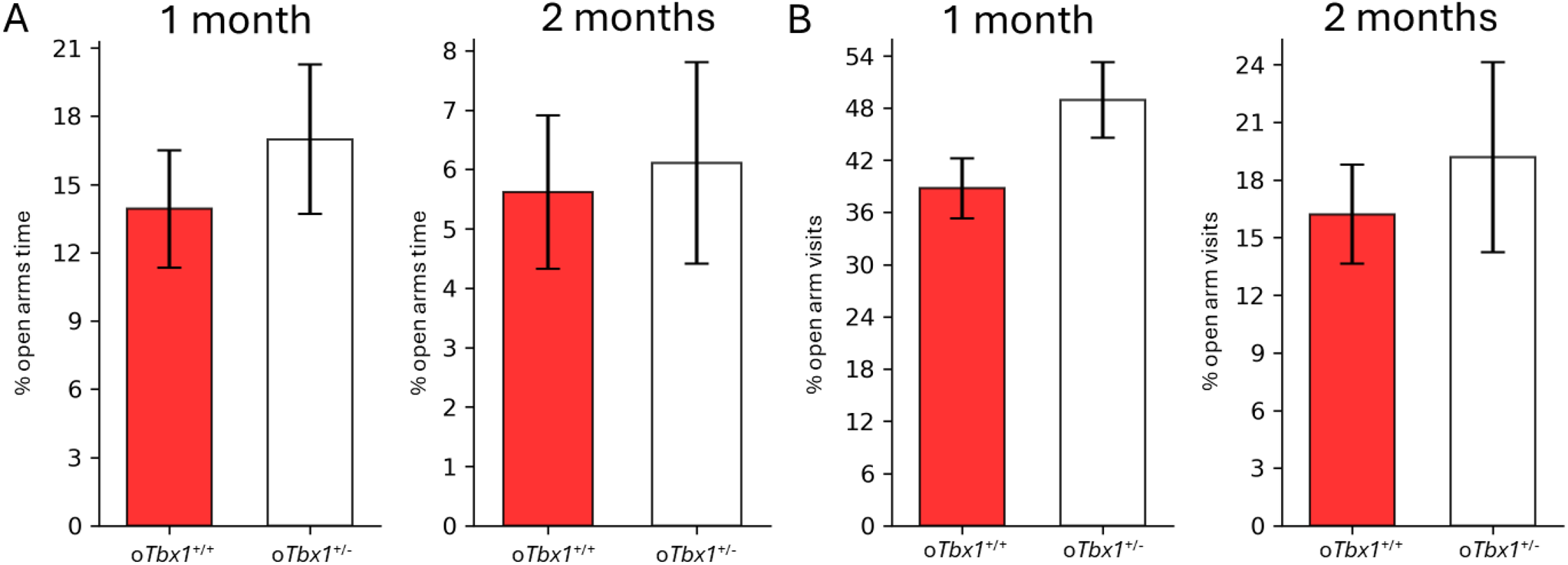
Anxiety-related behavior in an elevated T-maze.The *oTbx1*^+/−^ and *oTbx1*^+/+^ mice did not differ significantly in the relative amount of time spent (mean +SEM) in the open arms of an elevated plus maze (**A**,1 month,U = 100, p = 0.3440; 2 months, U = 98, p = 0.9483) or the frequency of visits to the open arms of the elevated plus maze (**B**, 1 month,t(32) = 1.7412, p = 0.0912; 2 months, t(28) = 0.5925, p = 0.5583). The three genotypes of *oTbx1*+/+ (1 month, N=7, PdgfrαCre;*Tbx1*^+/+^, N=7, Wild-type;*Tbx1*^+/+^, N=6, Wild-type;*Tbx1*+/flox; 2 months, N=8, PdgfrαCre;*Tbx1*^+/+^, N=6, Wild-type;*Tbx1*^+/+^, N=6, Wild-type;*Tbx1*^+/flox^) did not differ in the percentage of time spent in open arms (1 month,genotype, F(2, 20) = 0.0745, p = 0.9285; 2 months,genotype, F(2, 17) = 2.126, p = 0.1500) or the percentage of visits to open arms (1 month,genotype, F(2, 20) = 0.1837, p = 0.8336; 2 months,genotype, F(2, 17) = 1.4821, p = 0.2551), and thus were combined as *oTbx1*^+/+^. The sample sizes were: *oTbx1*^+/+^ mice, N = 23; *oTbx1*^+/−^ mice, N = 11 at 1 month; *oTbx1*^+/+^ mice, N = 20; *oTbx1*^+/−^ mice, N = 10 at 2 months.

We assessed motor activity and anxiety-related behavior in a high-stress open-field task(58, 62) where there is no option to escape to a closed space. In this task, o*Tbx1*^+/−^ and o*Tbx1*^+/+^ mice showed no differences in distance traveled at both 1 month and 2 months of age (**Figure 8A,B**) or in time spent in the margin zone (**Figure 8C,D**).

**Figure 8.**
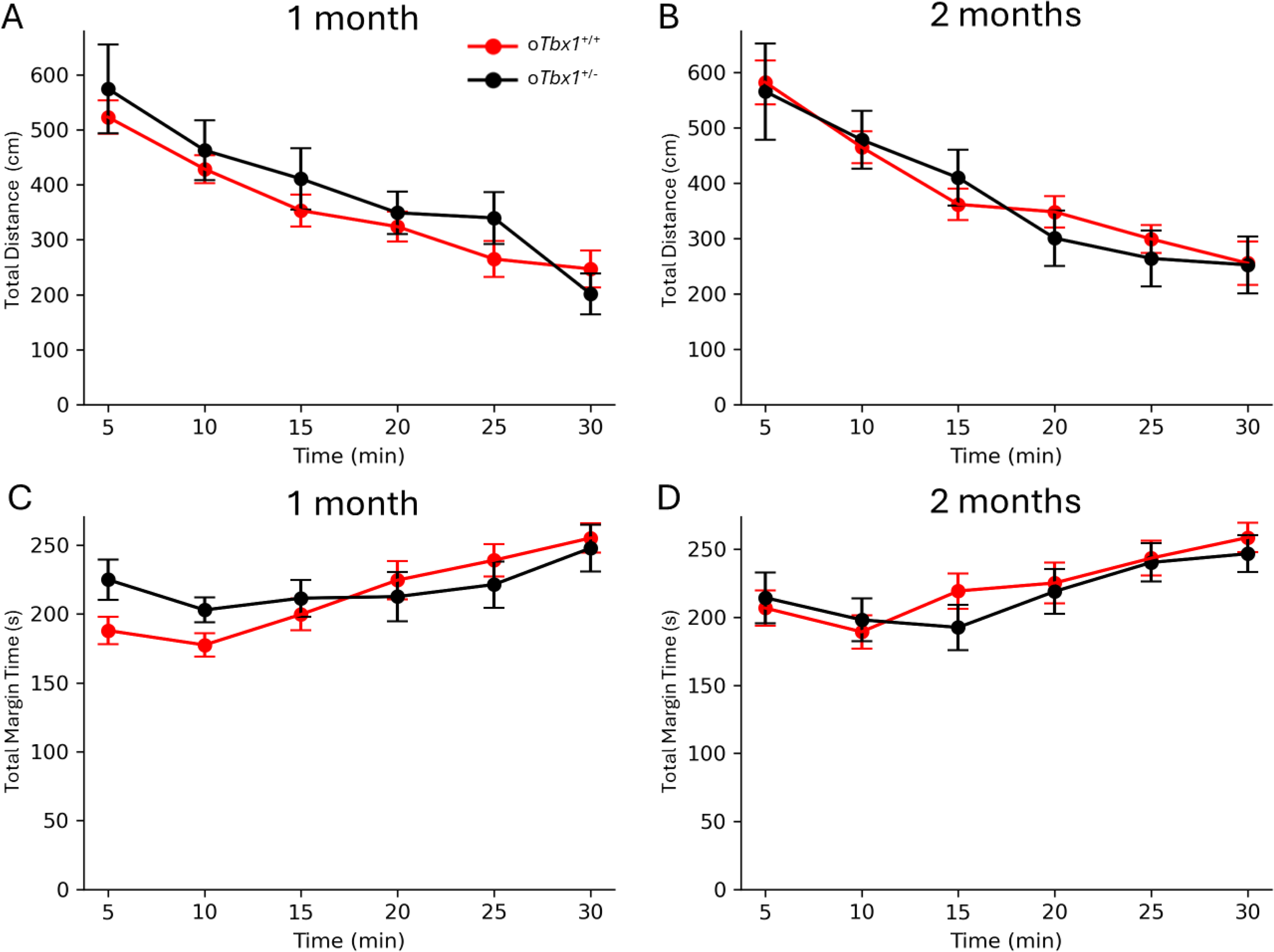
Motor Activity and Thigmotaxis. *oTbx1*^+/+^ and *oTbx1*^+/−^ mice exhibited indistinguishable distances traveled at both 1 month and 2 months of age (**A**, 1 month, genotype, F(1, 198) = 0.6314, p = 0.4278; genotype × time, F(5, 198) = 0.9551, p = 0.4466; **B**, 2 months, genotype, F(1, 28) = 0.0242, p = 0.87739; genotype × time, F(3.66, 102.485) = 0.5389, p = 0.69188). The Greenhouse–Geisser correction was applied due to a violation of sphericity, with the estimated epsilon being less than 0.75.The three genotypes of *oTbx1*^+/+^ (1 month, N=8, PdgfrαCre;*Tbx1*^+/+^, N=7, Wild-type;*Tbx1*^+/+^, N=9, Wild-type;*Tbx1*^+/flox^; 2 months, N=8, PdgfrαCre;*Tbx1*^+/+^, N=6, Wild-type;*Tbx1*^+/+^, N=6, Wild-type;*Tbx1*^+/flox^) demonstrated differences at both ages (genotype, F(2, 21) = 5.475, p = 0.01220; genotype × time, F(7.10, 74.58) = 1.118, p = 0.3611; at 2 months,genotype, F(2, 17) = 9.533, p = 0.00167; genotype × time, F(10, 85) = 0.3955, p = 0.9453). Comparisons of all pairs using Student’s t-tests indicated that no pair reached significance after the Benjamini-Hochberg correction (p>0.05 for all pairs), resulting in their combination as *oTbx1*^+/+^. *oTbx1*^+/+^ and *oTbx1*^+/−^ mice spent equal amounts of time in the margin zone (i.e., thigmotaxis) of the inescapable open field at both 1 and 2 months of age (**C,** 1 month, genotype, F(1, 198) = 0.2009, p = 0.6545; genotype × time, F(5,198) = 2.1517, p = 0.0609; **D**, 2 months, genotype, F(1, 168) = 0.08912, p = 0.7657; genotype × time, F(5, 168) = 0.9880, p = 0.4267). The three genotypes of *oTbx1*^+/+^ did not differ at 1 month of age (genotype, F(2, 21) = 0.3686, p = 0.6961; genotype × time, F(5.485, 57.59) = 0.9884, p = 0.4375), but differences were observed at 2 months of age (genotype, F(2, 17) = 4.880, p = 0.0211; genotype × time, F(5.049, 42.917) = 0.7180, p = 0.6147). Comparisons of all pairs using Student’s t-tests revealed that 2-month-old Wild-type;*Tbx1*^+/flox^ and Wild-type;*Tbx1*^+/+^ differed at the first four time points, even after the Benjamini-Hochberg correction (p<0.05). However, these groups had relatively small sample sizes (N=6 for both), and PdgfrαCre;*Tbx1*^+/+^and Wild-type;*Tbx1*^+/flox^ or PdgfrαCre;*Tbx1*^+/+^ and Wild-type;*Tbx1*^+/flox^ did not differ at any time point (p >0.05), thus they were combined as *oTbx1*^+/+^. Sample sizes included *oTbx1*^+/+^ mice, N = 24; *oTbx1*^+/−^ mice, N = 11 at 1 month; *oTbx1*^+/+^ mice, N = 20; *oTbx1*^+/−^ mice, N = 10 at 2 months.

While conditional *Tbx1* heterozygosity in the oligodendrocyte cell lineage resulted in higher spontaneous alternation scores at the longest inter-trial delay at one month of age, this phenotype contrasts with the impaired spontaneous alternation observed in constitutive *Tbx1* heterozygous mice (58, 62). It is noteworthy that *oTbx1*^+/+^ mice exhibited consistent decline in alternation rates to chance levels at a 30 s delay, whereas *oTbx1*^+/-^ mice did show this decline not from 15 s to 30 s delay (see **Figure 6A**, **1 month**), indicating a delayed decay of working memory or cognitive flexibility. Furthermore, this conditional *Tbx1* heterozygosity did not replicate the altered neonatal vocalizations, peri-adolescent or postnatal social interaction deficits, or heightened responses to novel, non-social objects or thigmotaxis observed in constitutive *Tbx1* heterozygous mice(58).

### Effects of *Tbx1* Heterozygosity in Oligodendrocyte Precursor Cells on Myelinated Axons in the Fimbria

Following behavioral testing, we examined myelinated axons in the fimbria utilizing electron microscopy. Constitutive *Tbx1* heterozygosity distinctly results in an increased proportion of myelinated axons with diameters ranging from 200 to 600 nm diameter, while concurrently reducing the proportion of axons measuring ≥700 nm and <1,200 nm. Additionally, axons within the 700–1,500 nm range demonstrated a thicker myelin sheath, with no larger myelinated axons identified in constitutive *Tbx1* heterozygous mice (62). We evaluated whether the ultrastructural alterations observed in the fimbria of constitutive *Tbx1*^+/−^ mice were mirrored in o*Tbx1*^+/−^ mice.

Densely packed axons were identified in the fimbria (**Figure 9A**). To evaluate relative myelin thickness, we analyzed the ratio of inner axon diameter to outer fiber diameter (i.e., the g-ratio) (**Figure 9B**). Similar to the findings in constitutive *Tbx1* mice, the g-ratio plateaued at approximately 0.8, which is the optimal ratio for signal conduction in the brain (88). However, no significant differences in g-ratios were noted between o*Tbx1*^+/−^ and o*Tbx1*^+/+^ mice across the full range of axon diameters.

**Figure 9.**
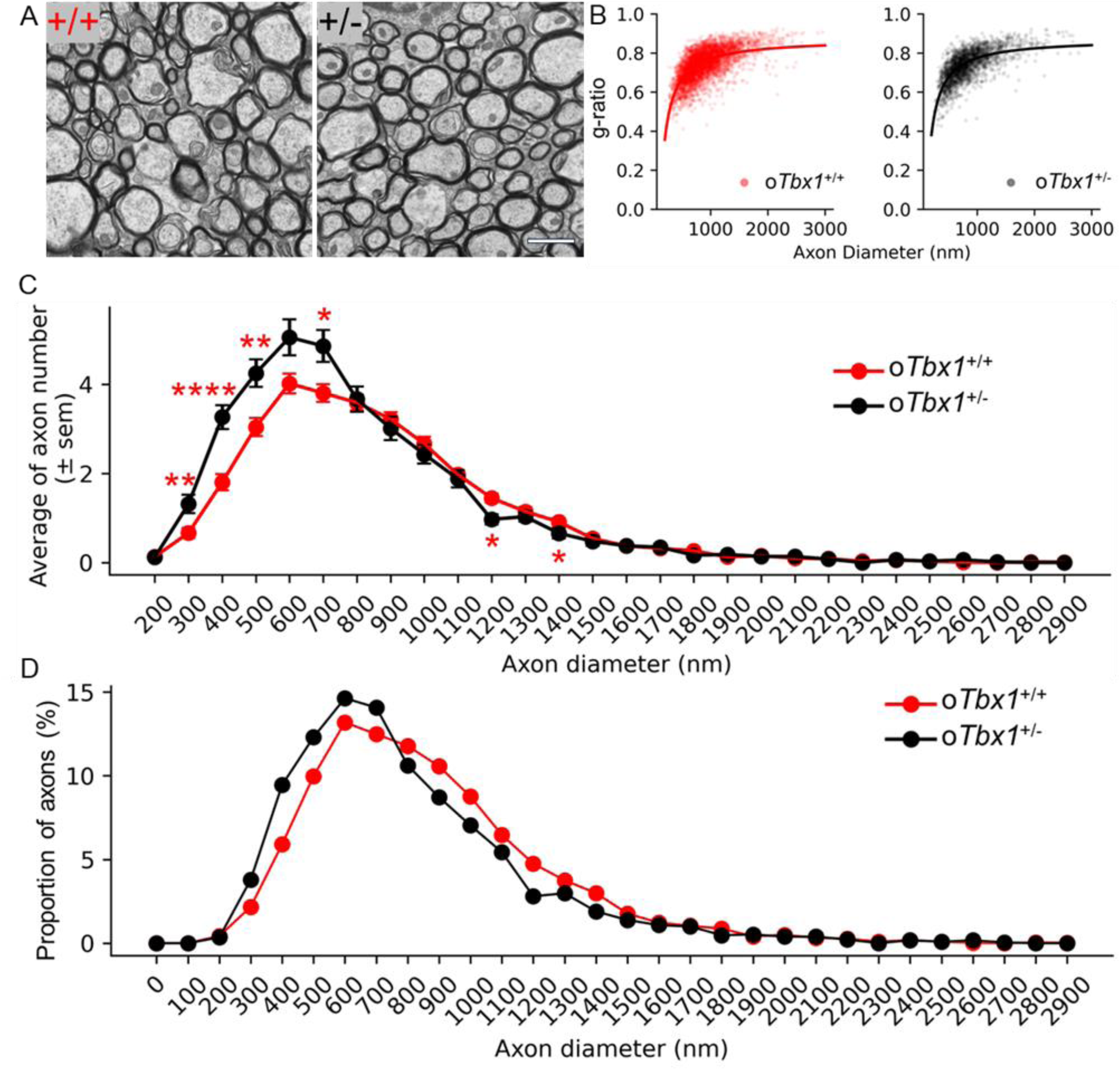
Electron Microscopy Analysis of Myelinated Axons. **A**. Representative electron microscopy images of axons in the fimbria at 20,000× magnification. Genotypes: +/+, *oTbx1*^+/+^; +/-, *oTbx1*^+/−^. Scale bar = 1,600 nm. **B**. G-ratios plotted against axon diameters for 4,423 and 2,317 axons from 5 *oTbx1*^+/+^ mice and 3 *oTbx1*^+/−^ mice, respectively. All axons with closed myelin sheaths were analyzed using MyelTracer. The reciprocal function best fits the data distributions. G-ratios increased as a function of axon diameter (*oTbx1*+/+, R = 0.6268, p < 0.001; *oTbx1*+/−, R = 0.6491, p < 0.001). There was no significant difference in the axon diameter range for G-ratios between oTbx1+/+ and oTbx1+/− mice (genotype, p = 0.8131; genotype × axon diameter, p = 0.221). **C**. Average number (± SEM) of axons per image field versus axon diameter. Significant differences were observed between *oTbx1*^+/+^ and *oTbx1*^+/−^ mice (χ² (27) = 109.7085, p = 6.22 × 10⁻¹²), primarily due to a higher number of axons with diameters ≥300 nm and <800 nm and a lower number of axons with diameters ≥1,200 nm and <1,500 nm in *oTbx1*^+/−^ compared to *oTbx1*^+/+^ mice, as determined by Mann-Whitney U tests applied to this diameter range (*, p < 0.05; **, p < 0.01; ***, p < 0.001; ****, p < 0.0001). The shift in axon sizes toward smaller to medium diameters in *oTbx1*^+/−^ compared to *oTbx1*^+/+^ mice resulted in a smaller average axon size for *oTbx1*^+/−^ mice (Mann-Whitney non-parametric tests, p = 7.48 × 10⁻¹⁹). **D**. Differences in the percentage of myelinated axons for *oTbx1*^+/−^ versus oTbx1+/+ mice across axon diameters (Kolmogorov-Smirnov, p = 3.59 × 10⁻¹⁶).

We also quantified the number and proportion of axons across different diameters. The fimbria of o*Tbx1*^+/−^ mice exhibited an increased number of axons measuring ≥ 300 nm and <800 nm, alongside decreased number of axons ≥1,200 nm and <1,500 nm diameters in comparison to o*Tbx1*^+/+^ mice (**Figure 9C**). Consequently, the curve representing relative axon proportions (y-axis) versus axon diameter (x-axis) shifted to the left in o*Tbx1*^+/−^ mice relative to o*Tbx1*^+/+^ mice (**Figure 9D**). Consistent with g-ratio findings, myelin thickness showed no significant differences between o*Tbx1*^+/−^ and *oTbx1*^+/+^ mice across all axon diameters (**Figure S2**). Thus, conditional *Tbx1* heterozygosity initiated in oligodendrocyte precursor cells selectively shifted the relative proportions of small to medium myelinated axons in the fimbria without impacting myelin thickness.

In summary, conditional *Tbx1* heterozygosity within the oligodendrocyte lineage resulted in a shift toward smaller-sized myelinated axons while leaving myelin thickness unchanged in the fimbria.

## DISCUSSION

The objective of this study was to delineate the cellular origin of the diverse behavioral and myelin phenotypes observed in constitutive *Tbx1* heterozygous mice(58-62). Given that social and cognitive phenotypes are recapitulated by initiating *Tbx1* heterozygosity in neonatal progenitor cells (72), and that certain neonatal neural progenitor cells in the mouse brain are responsible for generating neurons and oligodendrocytes during the neonatal period (70, 71, 108), it is possible that *Tbx1* deficiency in the neuronal or oligodendrocyte lineage contributes to the observed behavioral phenotypes. This study aimed to investigate the impact of *Tbx1* heterozygosity in the oligodendrocyte lineage on the behavioral and ultrastructural phenotypes.

To achieve this aim, we crossed *Pdgfrα*Cre mice with congenic *Tbx1*^+/flox^ mice to generate *Pdgfrα*Cre;*Tbx1*^+/flox^ (o*Tbx1*^+/-^) mice. We evaluated a range of social, cognitive, anxiety-related, and motor behaviors, as well as the ultrastructural composition of myelin and axons in the fimbria. The o*Tbx1*^+/-^ mice selectively replicated the altered proportion of axon sizes in the fimbria observed in constitutive *Tbx1* heterozygous mice. In contrast, o*Tbx1*^+/-^ exhibited higher levels of spontaneous alternation following a 30-second delay at one month of age—indicative of a delayed decay of working memory and cognitive flexibility-- contrary to the impaired spontaneous alternation observed in constitutive *Tbx1* heterozygous mice. This observation supports the notion that the final phenotype arises from the cumulative effects of various contributory and opposing factors (54, 80).

Other phenotypes associated with constitutive *Tbx1* heterozygosity were absent in o*Tbx1*^+/-^mice, including neonatal social communication, responses to social or non-social stimuli, and anxiety-like behaviors in an elevated plus maze and thigmotaxis. These findings indicate that *Tbx1* in the oligodendrocyte lineage does not significantly influence these phenotypes. Combined with the observation that *Tbx1* heterozygosity initiated in neonatal (P1–P5) neural progenitor cells recapitulates some social and cognitive deficits observed in constitutive *Tbx1* heterozygous mice (72), the current negative data suggest that *Tbx1* in neonatal neural progenitor cells or their neuronal lineage may play a principal role in determining the final phenotype (72).

We cannot dismiss the possibility that conditional *Tbx1* heterozygosity was incomplete. The *Pdgfrα* promoter is expected to induce nearly complete recombination in oligodendrocyte precursor cells, with labels persisting in their mature progeny, oligodendrocytes (109, 110); however, variability in recombination may occur in certain cells (111). The apparent absence of behavioral impairments may be attributed to incomplete recombination in all or some individual oligodendrocyte precursor cells. Nonetheless, o*Tbx1*^+/-^ mice showed an axonal phenotype at a presumed gene dosage reduction that did not significantly affect behavior, clearly indicating that behavioral phenotypes exhibit lower sensitivity to *Tbx1* heterozygosity in oligodendrocyte precursor cells compared to the axonal phenotype.

Constitutive *Tbx1* heterozygous mice exhibit increased myelin thickness of medium-sized axons and a lack of large (≥1,700 nm) myelinated axons in the fimbria(62); however, these characteristics were absent in o*Tbx1*^+/-^ mice. On the other hand, the fimbria of o*Tbx1*^+/-^ mice exhibited a greater number of axons with diameters ranging from ≥300 nm to <800 nm and a reduced number of axons with diameters from ≥1,200 nm to <1,500 nm, compared to o*Tbx1*^+/+^ mice. This pattern partially aligns with the increase in axons measuring ≥200 nm and <400 nm, as well as the decrease in axons measuring ≥700 nm and <1,700 nm in constitutive *Tbx1* heterozygous mice. The slight differences in the ranges of diameters of altered axons might be due to manual measurements in our previous study (62) and an automated measurements by MyelTracer in the current study.

This shift in axonal composition may result from oligodendrocyte lineage effects in a non-cell-autonomous manner, or from a non-specific, cell-autonomous effect of the *Pdgfrα* promoter in neurons. While *Pdgfrα*Cre primarily induces recombination in oligodendrocyte precursor cells, it also initiates recombination in neurons, astrocytes, pericytes, ependyma, perivascular mesenchymal cells, and other cell types(74-77, 99, 110, 112).

The recombination initiated by *Pdgfra*Cre could have influenced the behavioral phenotype in spontaneous alternation through interactions with other cell types and their progeny. In the mouse brain, *Tbx1* is undetectable in astrocytes or microglia, yet it is expressed in endothelial cells and neural progenitor cells. Furthermore, *Tbx1* is detected in immature neurons as well as both excitatory and inhibitory neurons at various developmental stages, from embryonic development to adulthood(65, 66, 113, 114). Additional research is required to clarify the impacts of *Tbx1* deficiency across different cell types on phenotypic outcomes.

The general absence of phenotypic abnormalities may be attributed to the high mortality rate observed in o*Tbx1*^+/-^ litters, potentially skewing our sample toward mice that were less affected by, or more resilient to, the mutation’s effects. In our breeder colony, only 31% and 24% of male pups survived to postnatal day (P7) and 2 months of age, respectively. Nevertheless, two lines of evidence do not support this explanation. First, a greater number of mice were alive during the neonatal period compared to 1 and 2 months of age, yet neonatal vocalizations on P8 and P12 were normal. Second, our nestinCreERT;*Tbx1*^+/flox^ mice also exhibited a high mortality rate but exhibited more significant behavioral deficits (72).

Large-scale human brain imaging studies have identified alterations in white matter among carriers of CNV(115, 116), as well as in idiopathic cases of ASD and schizophrenia (117, 118). Moreover, individuals with hemizygous deletions of 22q11.2 exhibit changes in white matter within the fornix/fimbria(119). Our present findings do not support the hypothesis that *Tbx1* deficiency in the oligodendrocyte lineage contributes to the structural or behavioral abnormalities related to 22q11.2 hemizygosity and protein-truncating variants of TBX1. Further research is necessary to identify the specific cell types responsible for the behavioral and structural phenotypes linked to *Tbx1* heterozygosity, 22q11.2 hemizygosity, and other CNVs, in order to establish a generalizable mechanistic basis for dimensions associated with mental illness.

---

**Table S1.**
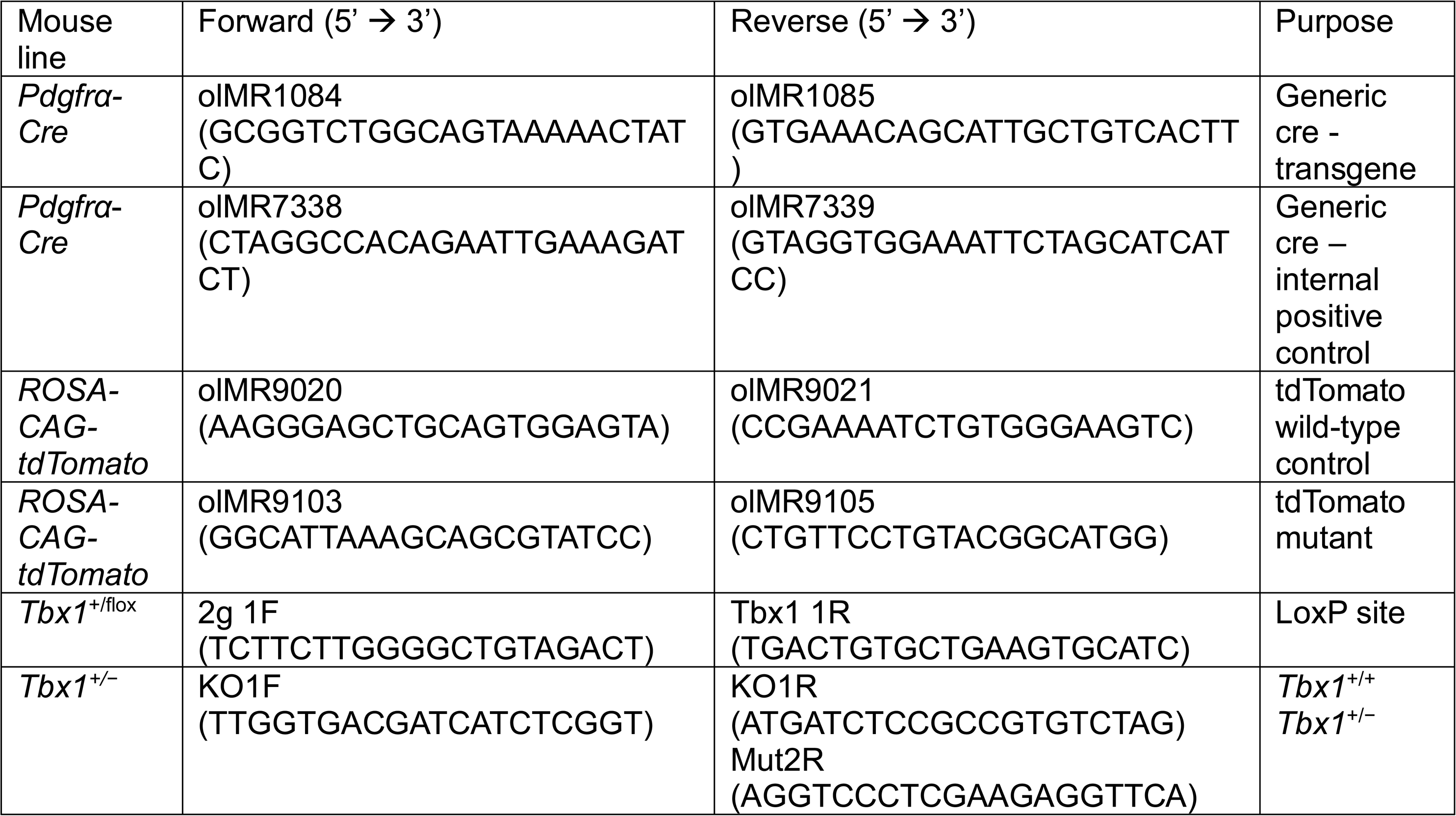
Primers used for genotyping.

**Table S2.**
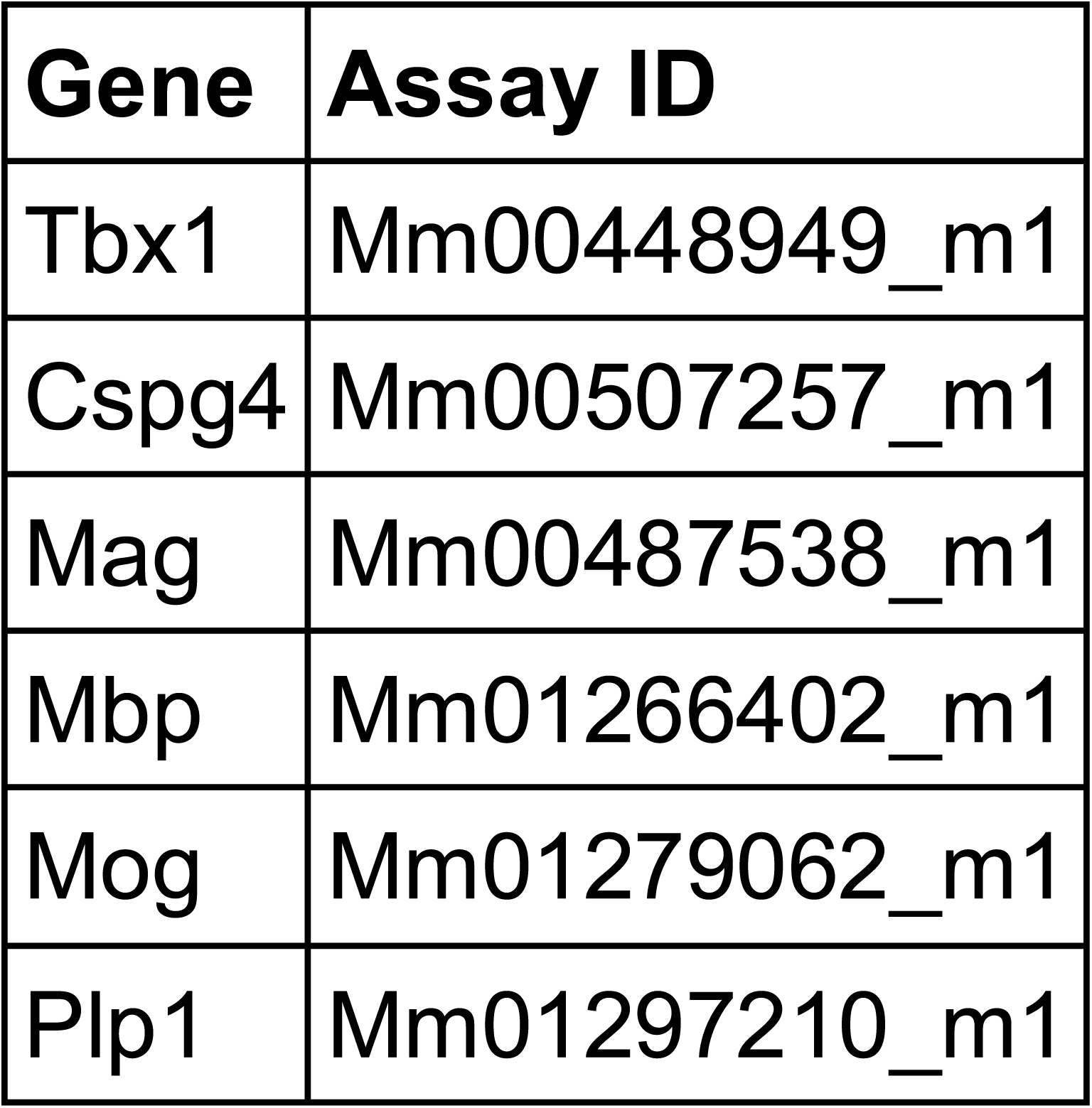
Primers for qRT-PCR.

| Gene | Assay ID |
| --- | --- |
| Tbx1 | Mm00448949_m1 |
| Cspg4 | Mm00507257_m1 |
| Mag | Mm00487538_m1 |
| Mbp | Mm01266402_m1 |
| Mog | Mm01279062_m1 |
| Plp1 | Mm01297210_m1 |

## Supplementary Information

**Figure S1.**
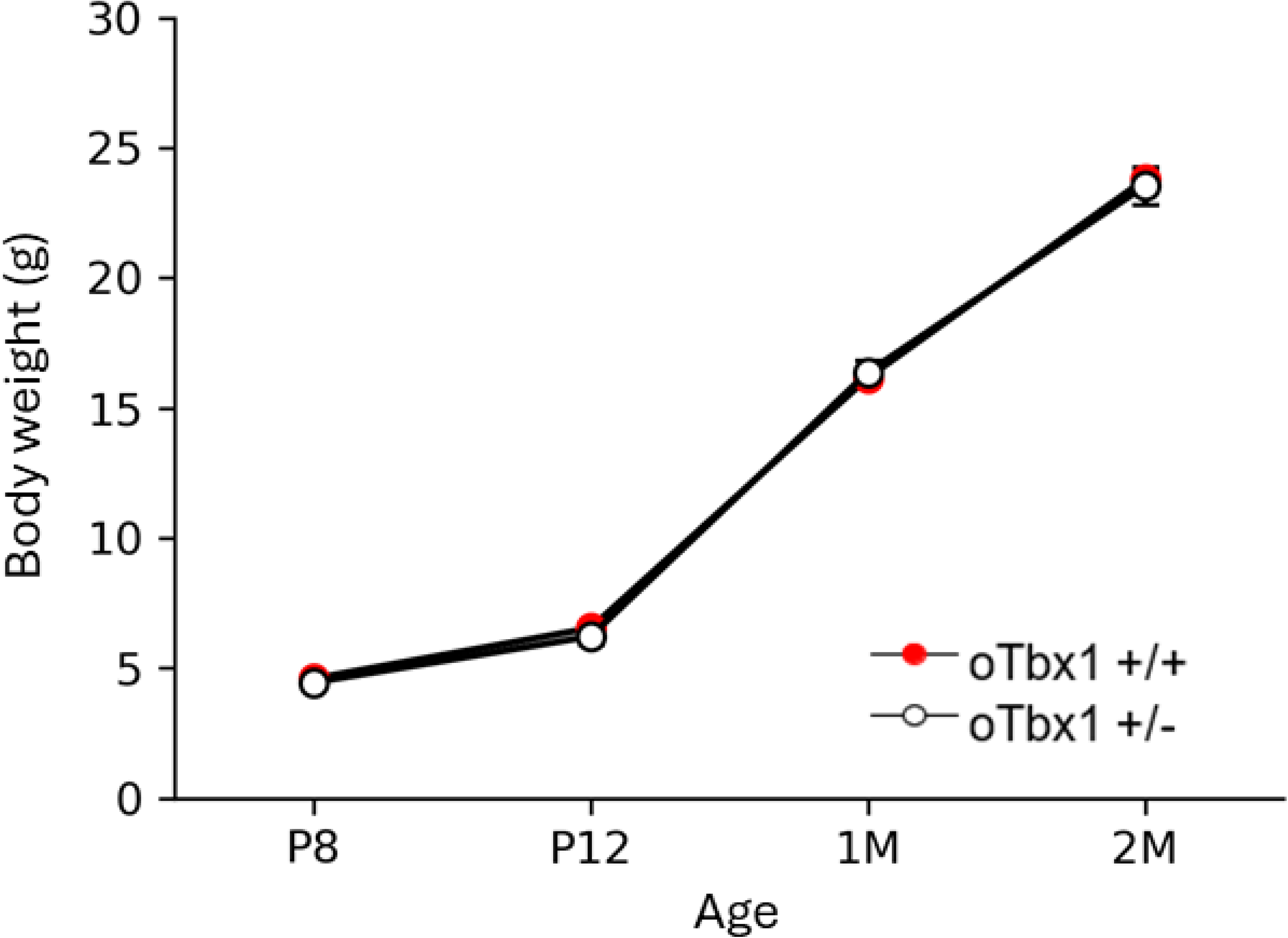
Body weights o*Tbx1*^+/+^ and o*Tbx1*^+/−^ mice. o*Tbx1*^+/+^ and o*Tbx1*^+/−^ mice had indistinguishable body weight increases from P8 to 2 months of age (P8, U = 140, p = 0.800; P12, U = 153, p = 0.466; 1 month, U = 135.5, p = 0.7035; 2 months, U = 102, p = 0.917). o*Tbx1*^+/+^ P8, N = 27, P12; N = 30; 1M, 1 month, N = 27; 2M, 2 months, N = 21: o*Tbx1*^+/−^: P8, N = 11, P12, N = 12; 1M, N = 11; 2M, N = 10.

**Supplementary Figure S2.**
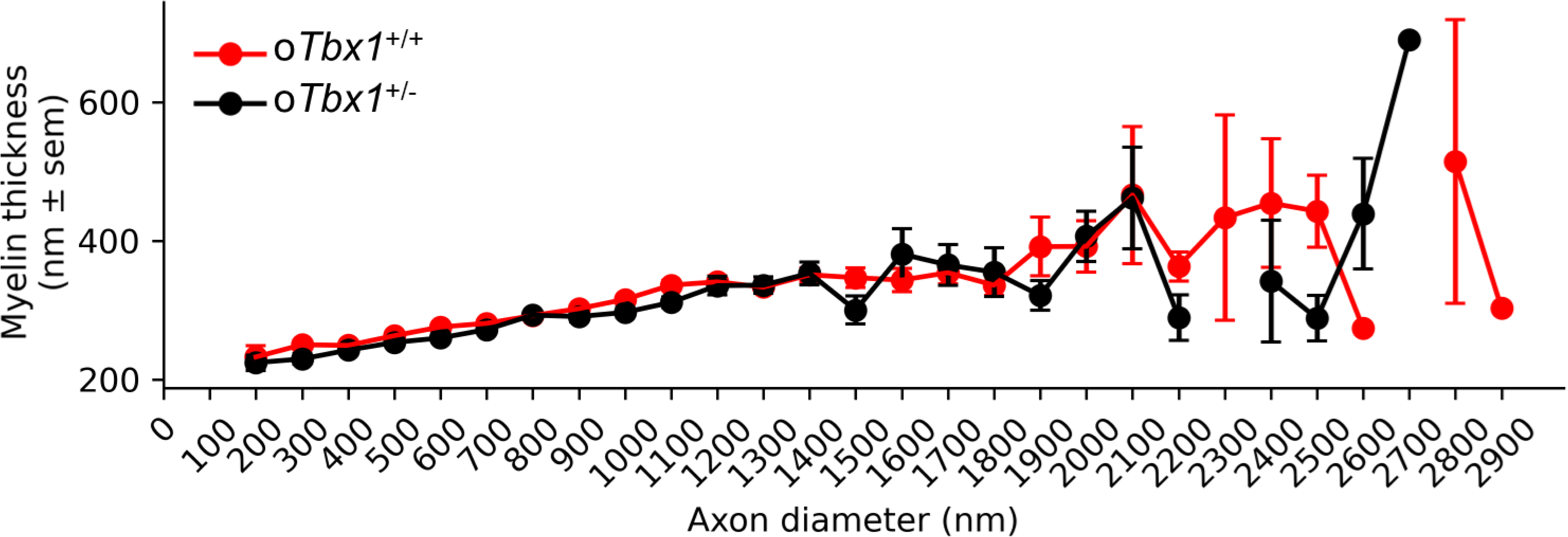
The myelin thickness of axons for o*Tbx1*^+/−^ versus o*Tbx1*^+/+^ mice versus axon diameter was similar along the axon diameter range (genotype, p = 0.931; genotype × axon diameter, p = 0.814).

---

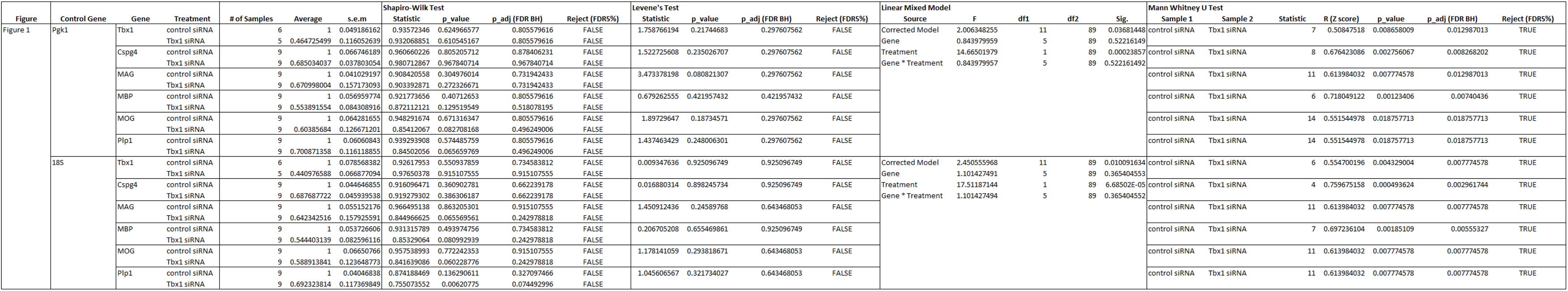

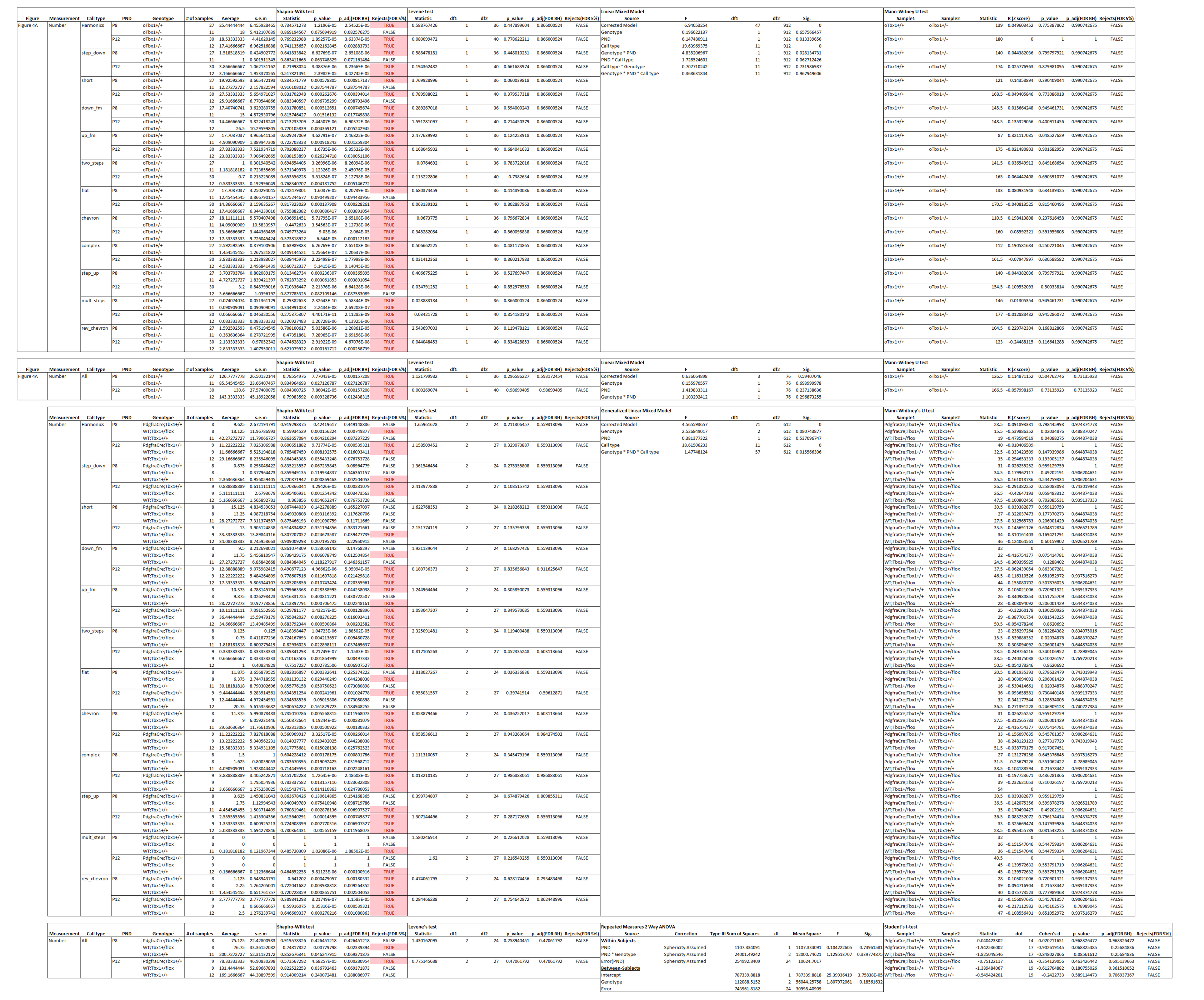

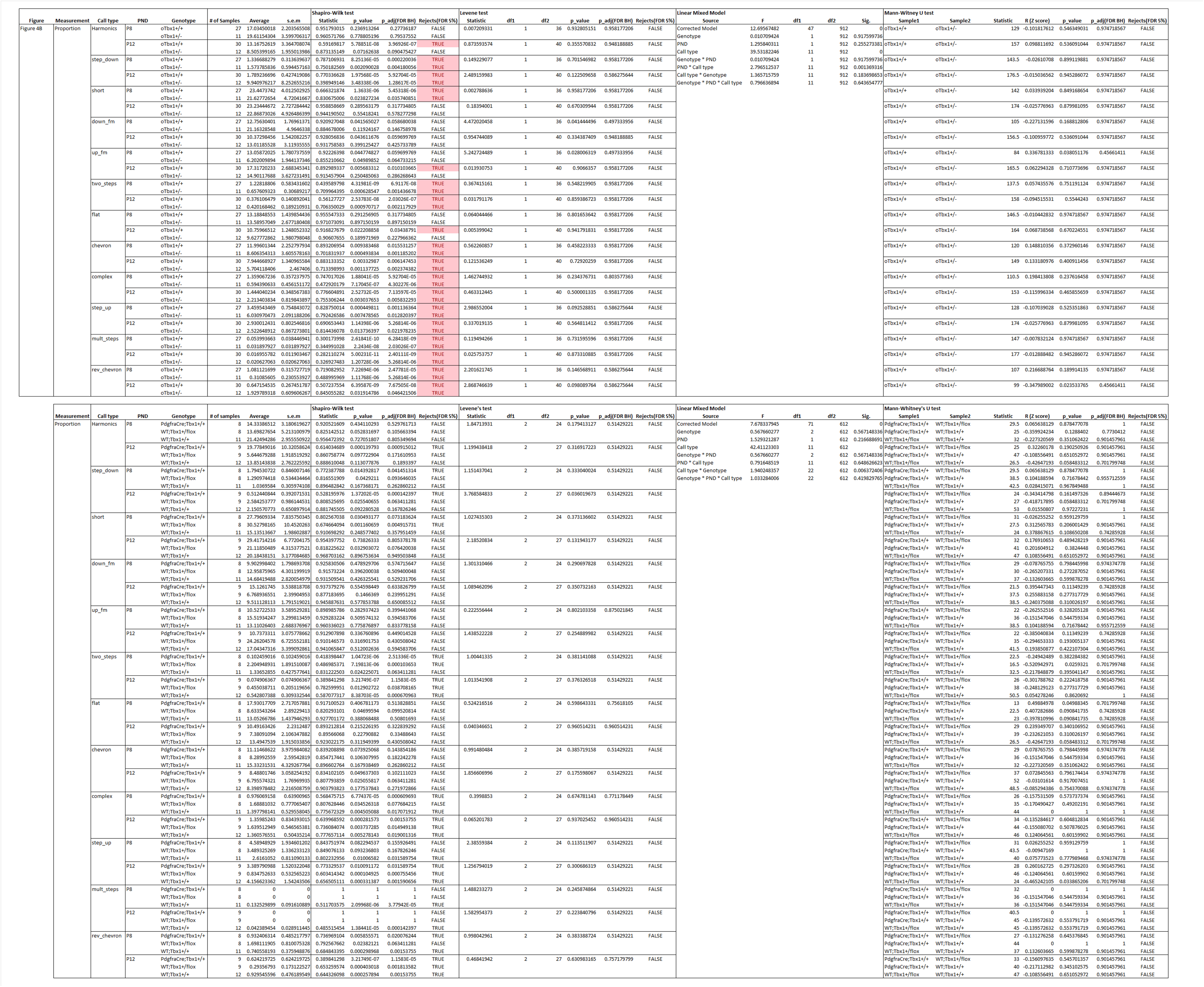

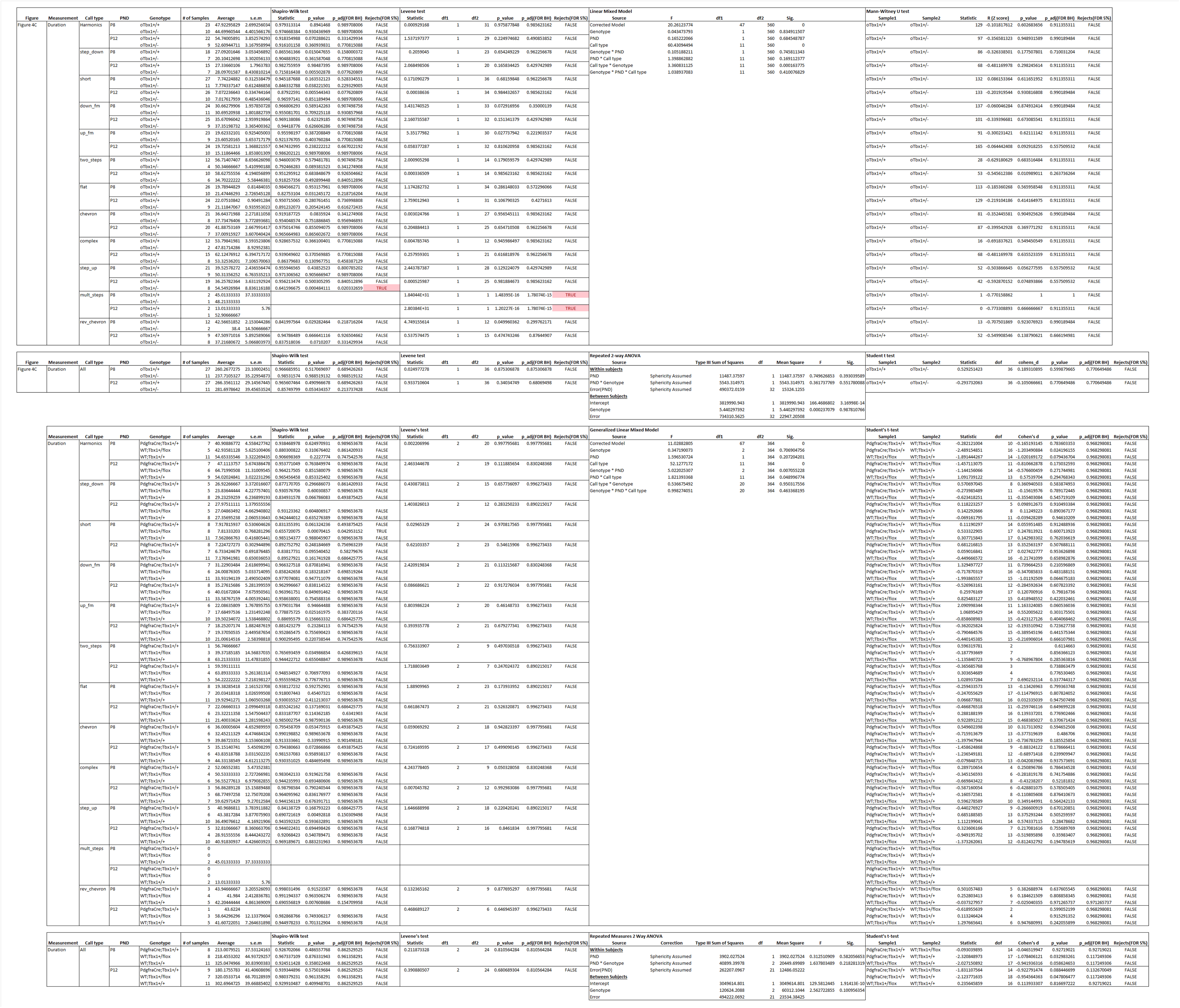

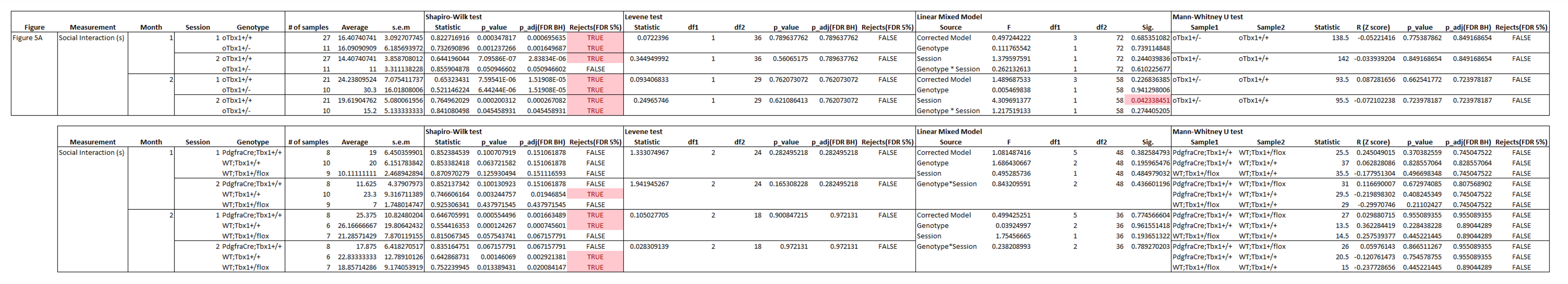

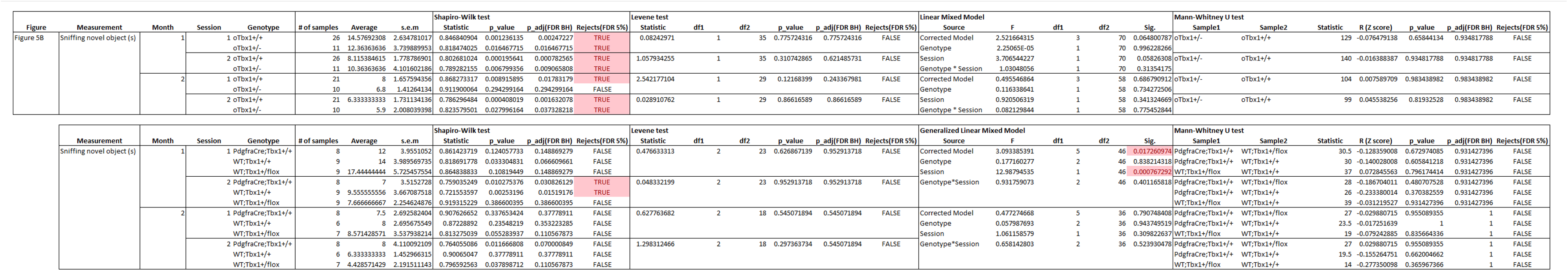

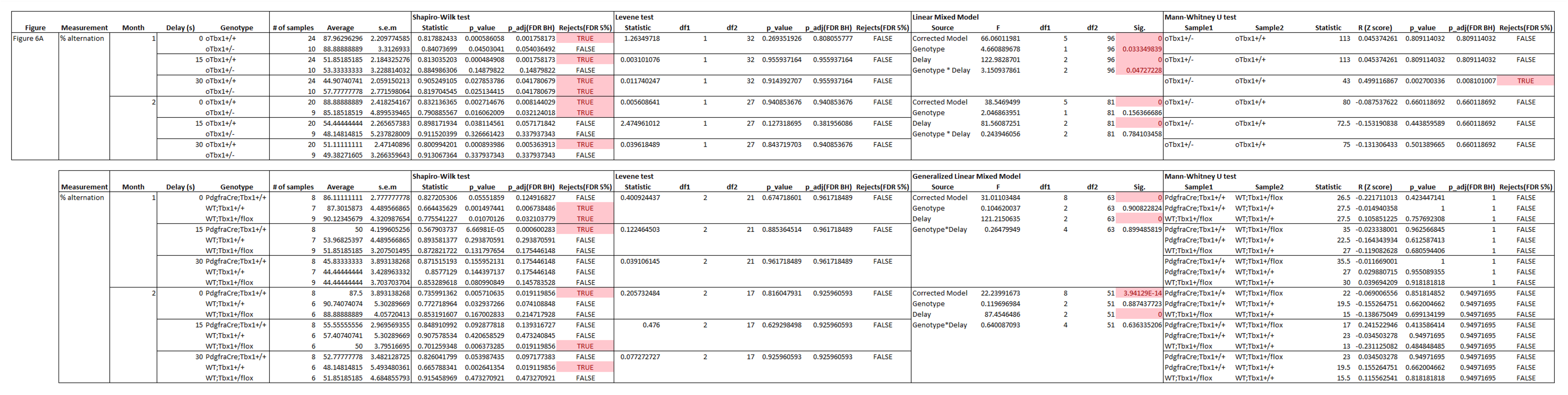

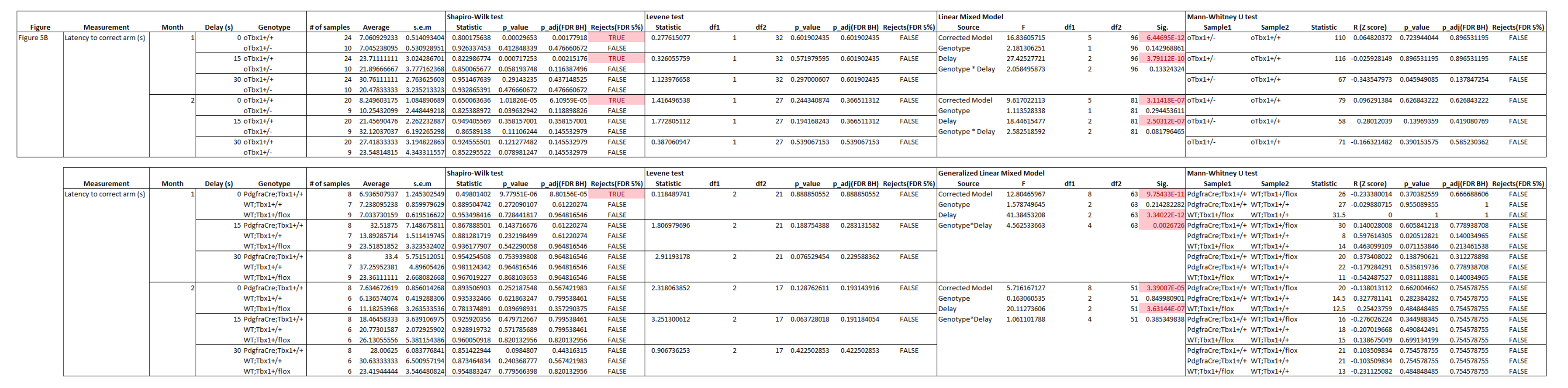

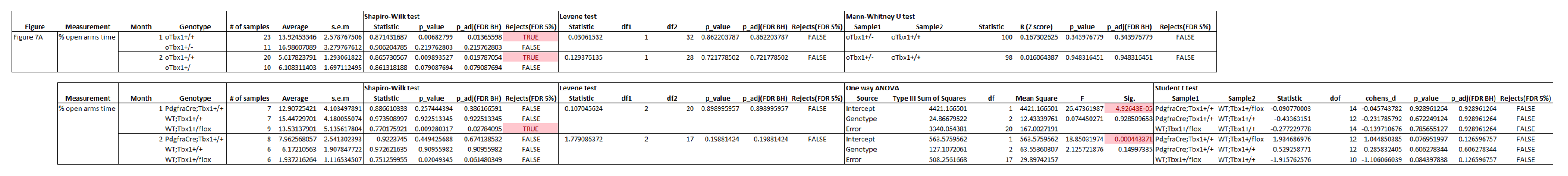

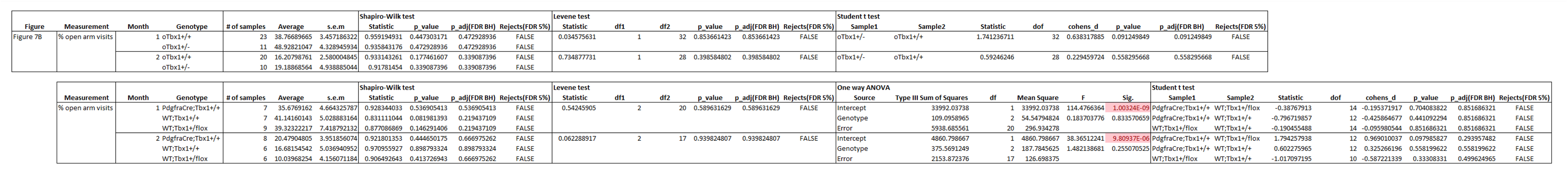

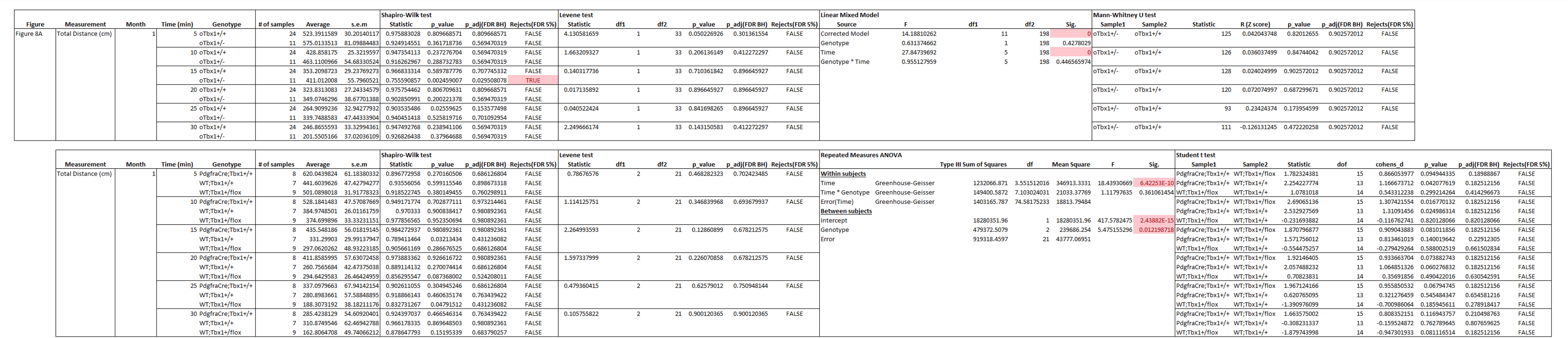

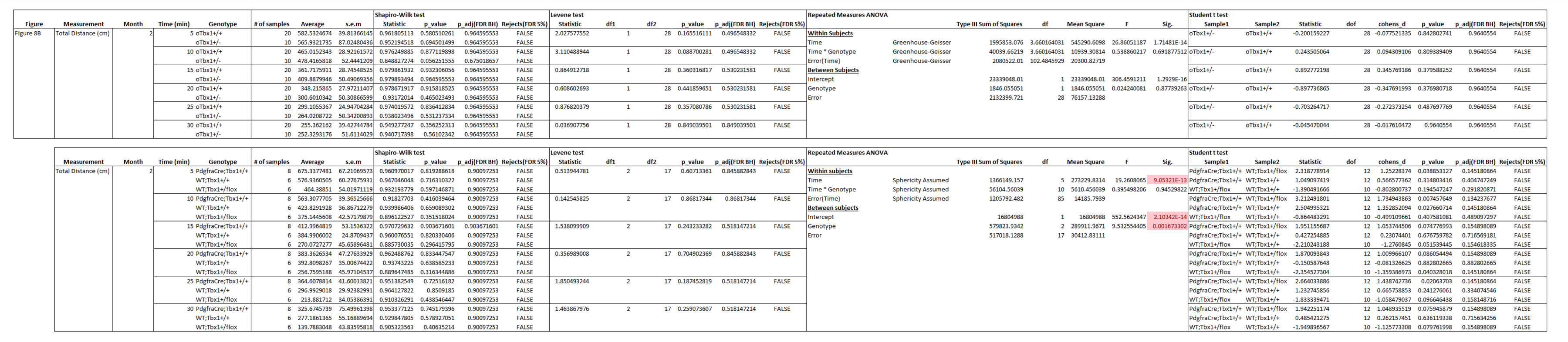

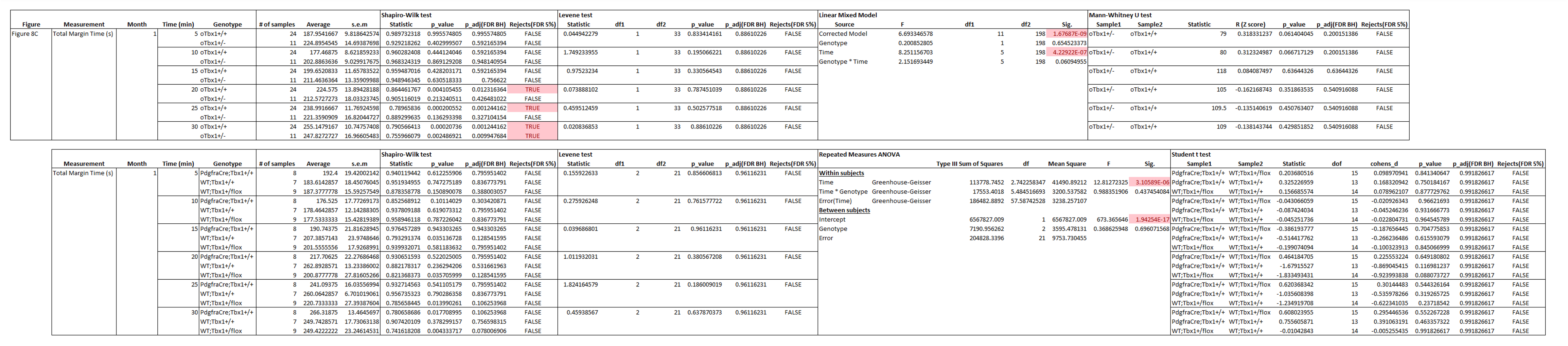

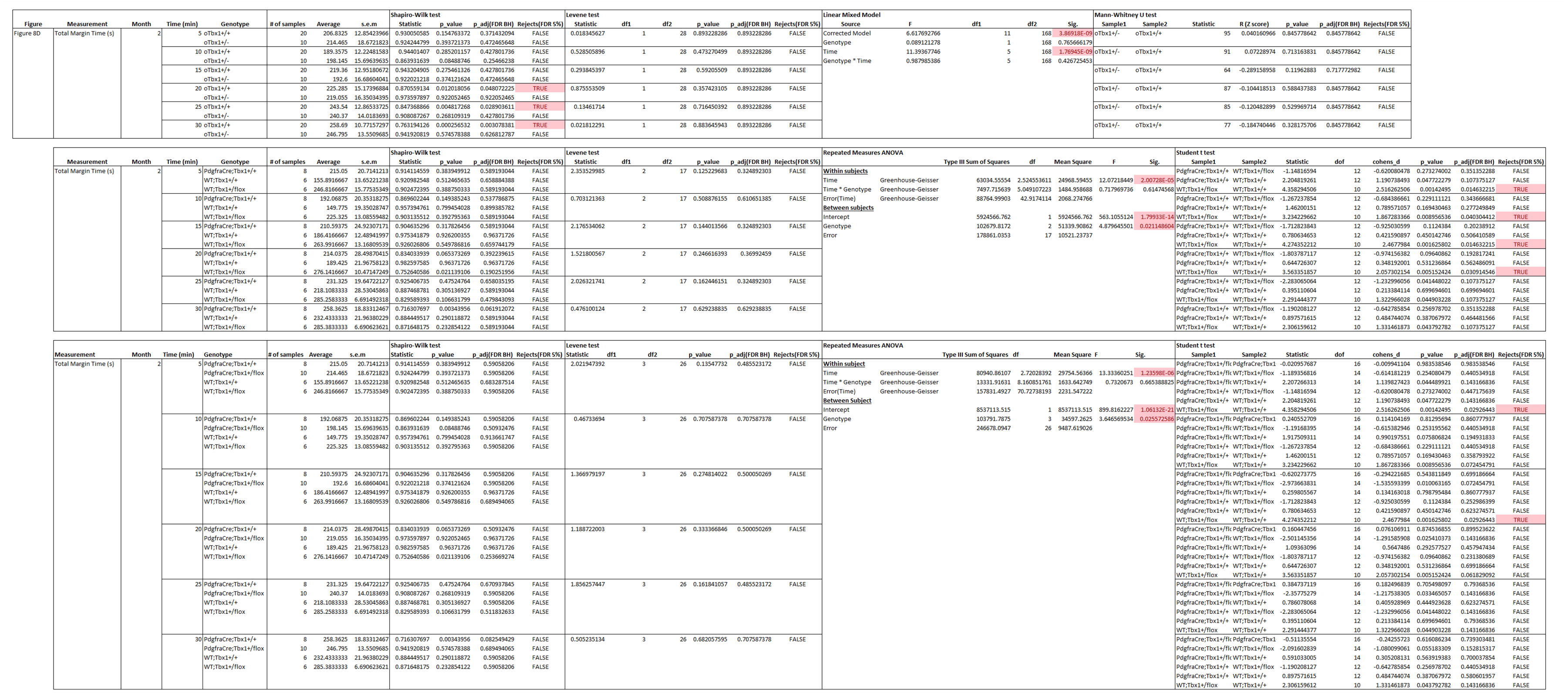

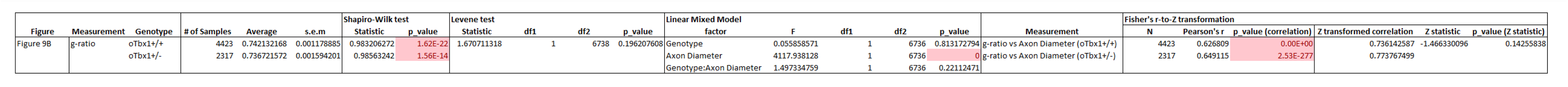

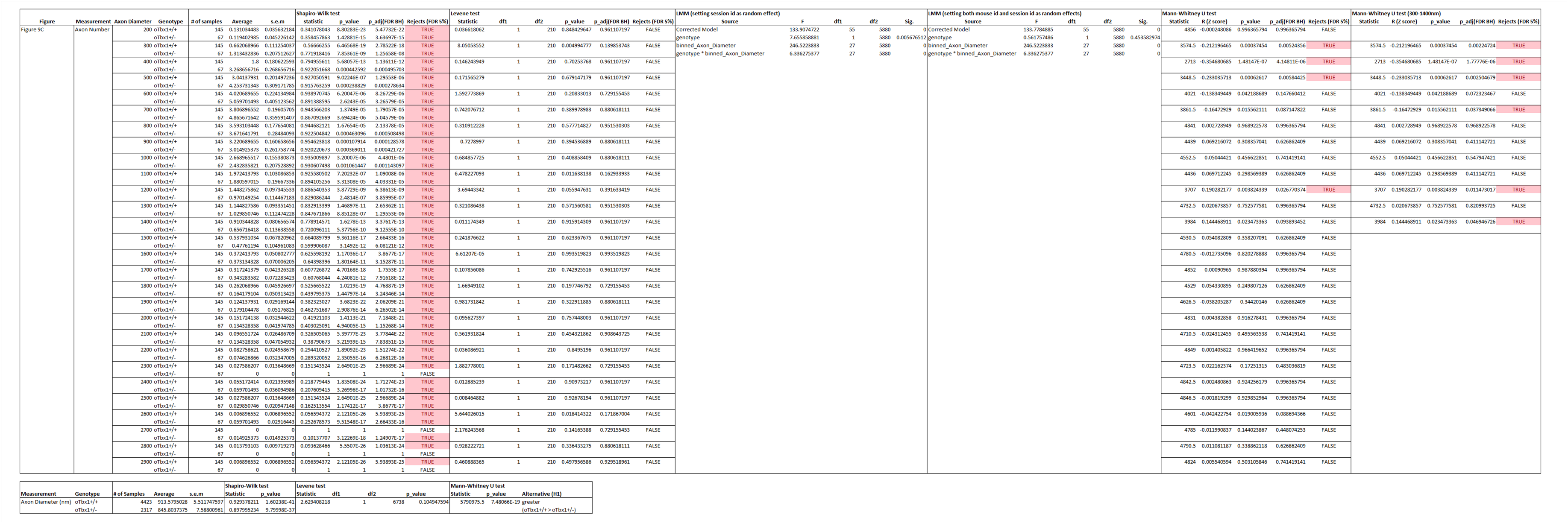

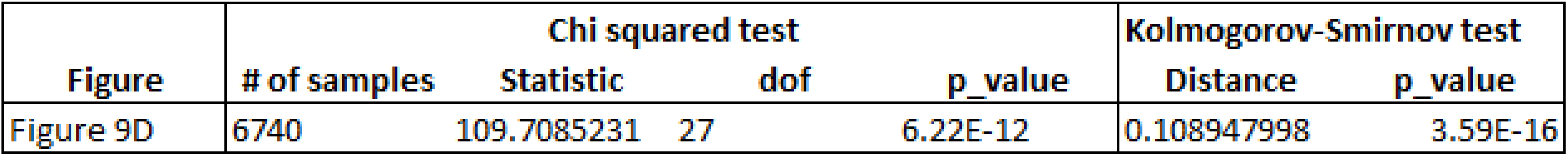

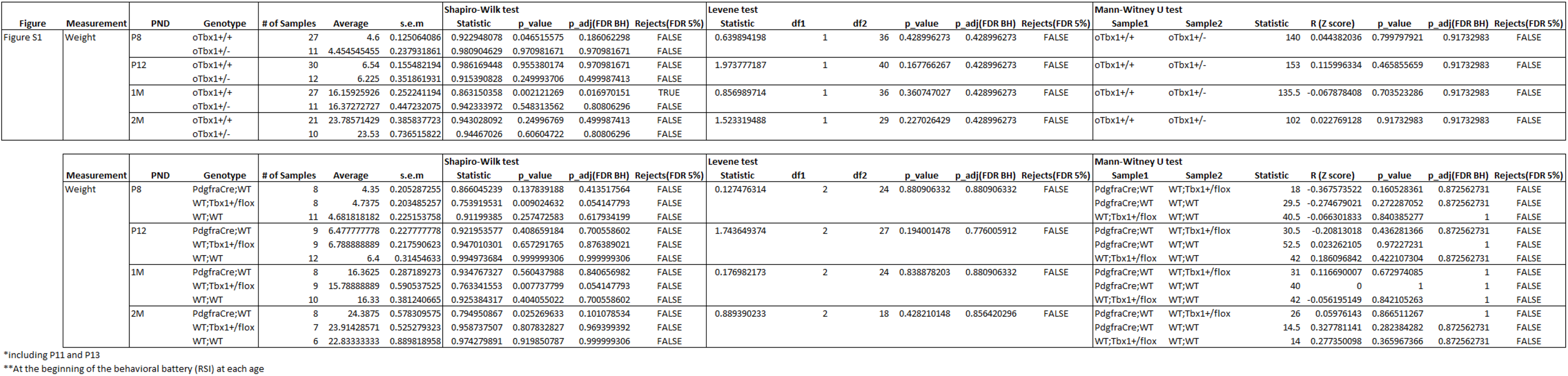

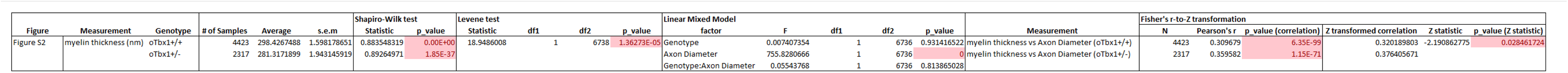

## Notes

### Competing Interest Statement

The authors have declared no competing interest.

### Summary of Updates

The text responded to reviewers' comments.

## References

1. Hiroi N, Yamauchi T. Modeling and Predicting Developmental Trajectories of Neuropsychiatric Dimensions Associated With Copy Number Variations. Int J Neuropsychopharmacol. 2019;22(8):488–500.

2. Michelini G, Carlisi CO, Eaton NR, Elison JT, Haltigan JD, Kotov R, et al. Where do neurodevelopmental conditions fit in transdiagnostic psychiatric frameworks? Incorporating a new neurodevelopmental spectrum. World Psychiatry. 2024;23(3):333–57.

3. Ozonoff S, Iosif AM, Baguio F, Cook IC, Hill MM, Hutman T, et al. A prospective study of the emergence of early behavioral signs of autism. J Am Acad Child Adolesc Psychiatry. 2010;49(3):256–66.

4. Ozonoff S, Young GS, Belding A, Hill M, Hill A, Hutman T, et al. The broader autism phenotype in infancy: when does it emerge? J Am Acad Child Adolesc Psychiatry. 2014;53(4):398–407.

5. Esposito G, Hiroi N, Scattoni ML. Cry, baby, cry: Expression of Distress as a Biomarker and Modulator in Autism Spectrum Disorder. Int J Neuropsychopharmacol. 2017;20(6):498–503.

6. Sung S, Fenoglio A, Wolff JJ, Schultz RT, Botteron KN, Dager SR, et al. Examining the factor structure and discriminative utility of the Infant Behavior Questionnaire-Revised in infant siblings of autistic children. Child Dev. 2022;93(5):1398–413.

7. Northrup JB, Leezenbaum NB, Campbell SB. Observed Social Emotional Behavior at 22 Months Predicts a Later ASD Diagnosis in High-Risk Siblings. J Autism Dev Disord. 2021;51(9):3187–98.

8. Miller M, Sun S, Iosif AM, Young GS, Belding A, Tubbs A, et al. Repetitive behavior with objects in infants developing autism predicts diagnosis and later social behavior as early as 9 months. J Abnorm Psychol. 2021;130(6):665–75.

9. Di Giorgio E, Rosa-Salva O, Frasnelli E, Calcagni A, Lunghi M, Scattoni ML, et al. Abnormal visual attention to simple social stimuli in 4-month-old infants at high risk for Autism. Sci Rep. 2021;11(1):15785.

10. Miller M, Iosif AM, Hill M, Young GS, Schwichtenberg AJ, Ozonoff S. Response to Name in Infants Developing Autism Spectrum Disorder: A Prospective Study. J Pediatr. 2017;183:141–6 e1.

11. Caravella KE, Roberts JE. Adaptive Skill Trajectories in Infants with Fragile X Syndrome Contrasted to Typical Controls and Infants at High Risk for Autism. Res Autism Spectr Disord. 2017;40:1–12.

12. Gur RC, Calkins ME, Satterthwaite TD, Ruparel K, Bilker WB, Moore TM, et al. Neurocognitive growth charting in psychosis spectrum youths. JAMA Psychiatry. 2014;71(4):366–74.

13. Seidman LJ, Shapiro DI, Stone WS, Woodberry KA, Ronzio A, Cornblatt BA, et al. Association of Neurocognition With Transition to Psychosis: Baseline Functioning in the Second Phase of the North American Prodrome Longitudinal Study. JAMA Psychiatry. 2016;73(12):1239–48.

14. Meier MH, Caspi A, Reichenberg A, Keefe RS, Fisher HL, Harrington H, et al. Neuropsychological decline in schizophrenia from the premorbid to the postonset period: evidence from a population-representative longitudinal study. Am J Psychiatry. 2014;171(1):91–101.

15. Bora E, Lin A, Wood SJ, Yung AR, McGorry PD, Pantelis C. Cognitive deficits in youth with familial and clinical high risk to psychosis: a systematic review and meta-analysis. Acta Psychiatr Scand. 2014;130(1):1–15.

16. Sheffield JM, Karcher NR, Barch DM. Cognitive Deficits in Psychotic Disorders: A Lifespan Perspective. Neuropsychol Rev. 2018;28(4):509–33.

17. Tschentscher N, Woll CFJ, Tafelmaier JC, Kriesche D, Bucher JC, Engel RR, et al. Neurocognitive Deficits in First-Episode and Chronic Psychotic Disorders: A Systematic Review from 2009 to 2022. Brain Sci. 2023;13(2).

18. Johansson M, Hjarthag F, Helldin L. Cognitive markers related to long-term remission status in Schizophrenia Spectrum Disorders. Psychiatry Res. 2020;289:113035.

19. Stefansson H, Meyer-Lindenberg A, Steinberg S, Magnusdottir B, Morgen K, Arnarsdottir S, et al. CNVs conferring risk of autism or schizophrenia affect cognition in controls. Nature. 2014;505(7483):361–6.

20. Kendall KM, Rees E, Escott-Price V, Einon M, Thomas R, Hewitt J, et al. Cognitive Performance Among Carriers of Pathogenic Copy Number Variants: Analysis of 152,000 UK Biobank Subjects. Biol Psychiatry. 2017;82(2):103–10.

21. Malhotra D, Sebat J. CNVs: harbingers of a rare variant revolution in psychiatric genetics. Cell. 2012;148(6):1223–41.

22. Marshall CR, Howrigan DP, Merico D, Thiruvahindrapuram B, Wu W, Greer DS, et al. Contribution of copy number variants to schizophrenia from a genome-wide study of 41,321 subjects. Nat Genet. 2017;49(1):27–35.

23. Goh S, Thiyagarajan L, Dudding-Byth T, Mark P, Kirk EP. A systematic review and pooled analysis of penetrance estimates of copy number variants associated with neurodevelopment. Genet Med. 2024:101227.

24. Gur RE, Yi JJ, Donald-McGinn DM, Tang SX, Calkins ME, Whinna D, et al. Neurocognitive development in 22q11.2 deletion syndrome: comparison with youth having developmental delay and medical comorbidities. Mol Psychiatry. 2014;19(11):1205–11.

25. Morrison S, Chawner S, van Amelsvoort T, Swillen A, Vingerhoets C, Vergaelen E, et al. Cognitive deficits in childhood, adolescence and adulthood in 22q11.2 deletion syndrome and association with psychopathology. Transl Psychiatry. 2020;10(1):53.

26. Chawner S, Owen MJ, Holmans P, Raymond FL, Skuse D, Hall J, et al. Genotype-phenotype associations in children with copy number variants associated with high neuropsychiatric risk in the UK (IMAGINE-ID): a case-control cohort study. Lancet Psychiatry. 2019;6(6):493–505.

27. Zinkstok J, Boot E, Bassett AS, Hiroi N, Butcher NJ, Vingerhoets C, et al. The 22q11.2 deletion syndrome from a neurobiological perspective. Lancet Psychiatry. 2019;6(11):951–60.

28. Schneider M, Debbane M, Bassett AS, Chow EW, Fung WL, van den Bree M, et al. Psychiatric disorders from childhood to adulthood in 22q11.2 deletion syndrome: results from the International Consortium on Brain and Behavior in 22q11.2 Deletion Syndrome. Am J Psychiatry. 2014;171(6):627–39.

29. Ensenauer RE, Adeyinka A, Flynn HC, Michels VV, Lindor NM, Dawson DB, et al. Microduplication 22q11.2, an emerging syndrome: clinical, cytogenetic, and molecular analysis of thirteen patients. American Journal of Human Genetics. 2003;73(5):1027–40.

30. Hassed SJ, Hopcus-Niccum D, Zhang L, Li S, Mulvihill JJ. A new genomic duplication syndrome complementary to the velocardiofacial (22q11 deletion) syndrome. Clinical Genetics. 2004;65(5):400–4.

31. Portnoi MF, Lebas F, Gruchy N, Ardalan A, Biran-Mucignat V, Malan V, et al. 22q11.2 duplication syndrome: two new familial cases with some overlapping features with DiGeorge/velocardiofacial syndromes. American Journal of Medical Genetics A. 2005;137(1):47–51.

32. Yobb TM, Somerville MJ, Willatt L, Firth HV, Harrison K, MacKenzie J, et al. Microduplication and triplication of 22q11.2: a highly variable syndrome. American Journal of Human Genetics. 2005;76(5):865–76.

33. de La Rochebrochard C, Joly-Helas G, Goldenberg A, Durand I, Laquerriere A, Ickowicz V, et al. The intrafamilial variability of the 22q11.2 microduplication encompasses a spectrum from minor cognitive deficits to severe congenital anomalies. Am J Med Genet A. 2006;140(14):1608–13.

34. Alberti A, Romano C, Falco M, Cali F, Schinocca P, Galesi O, et al. 1.5 Mb de novo 22q11.21 microduplication in a patient with cognitive deficits and dysmorphic facial features. Clin Genet. 2007;71(2):177–82.

35. Engels H, Brockschmidt A, Hoischen A, Landwehr C, Bosse K, Walldorf C, et al. DNA microarray analysis identifies candidate regions and genes in unexplained mental retardation. Neurology. 2007;68(10):743–50.

36. Mukaddes NM, Herguner S. Autistic disorder and 22q11.2 duplication. World J Biol Psychiatry. 2007;8(2):127–30.

37. Torres-Juan L, Rosell J, Morla M, Vidal-Pou C, Garcia-Algas F, de la Fuente MA, et al. Mutations in TBX1 genocopy the 22q11.2 deletion and duplication syndromes: a new susceptibility factor for mental retardation. Eur J Hum Genet. 2007;15(6):658–63.

38. Descartes M, Franklin J, de Stahl TD, Piotrowski A, Bruder CE, Dumanski JP, et al. Distal 22q11.2 microduplication encompassing the BCR gene. Am J Med Genet A. 2008;146A(23):3075–81.

39. Ramelli GP, Silacci C, Ferrarini A, Cattaneo C, Visconti P, Pescia G. Microduplication 22q11.2 in a child with autism spectrum disorder: clinical and genetic study. Dev Med Child Neurol. 2008;50(12):953–5.

40. Wentzel C, Fernstrom M, Ohrner Y, Anneren G, Thuresson AC. Clinical variability of the 22q11.2 duplication syndrome. Eur J Med Genet. 2008;51(6):501–10.

41. Yu S, Cox K, Friend K, Smith S, Buchheim R, Bain S, et al. Familial 22q11.2 duplication: a three-generation family with a 3-Mb duplication and a familial 1.5-Mb duplication. Clin Genet. 2008;73(2):160–4.

42. Lo-Castro A, Galasso C, Cerminara C, El-Malhany N, Benedetti S, Nardone AM, et al. Association of syndromic mental retardation and autism with 22q11.2 duplication. Neuropediatrics. 2009;40(3):137–40.

43. Portnoi MF. Microduplication 22q11.2: a new chromosomal syndrome. Eur J Med Genet. 2009;52(2-3):88–93.

44. Soysal Y, Vermeesch J, Davani NA, Sensoy N, Hekimler K, Imirzalioglu N. Molecular characterization of microduplication 22q11.2 in a girl with hypernasal speech. Genet Mol Res. 2011;10(3):2148–54.

45. Wenger TL, Miller JS, DePolo LM, de Marchena AB, Clements CC, Emanuel BS, et al. 22q11.2 duplication syndrome: elevated rate of autism spectrum disorder and need for medical screening. Mol Autism. 2016;7:27.

46. Singh T, Poterba T, Curtis D, Akil H, Al Eissa M, Barchas JD, et al. Rare coding variants in ten genes confer substantial risk for schizophrenia. Nature. 2022;604(7906):509–16.

47. Alhazmi S, Alzahrani M, Farsi R, Alharbi M, Algothmi K, Alburae N, et al. Multiple Recurrent Copy Number Variations (CNVs) in Chromosome 22 Including 22q11.2 Associated with Autism Spectrum Disorder. Pharmgenomics Pers Med. 2022;15:705–20.

48. Alghamdi M, Al Khalifah R, Al Homyani DK, Alkhamis WH, Arold ST, Ekhzaimy A, et al. A Novel TBX1 Variant Causing Hypoparathyroidism and Deafness. J Endocr Soc. 2020;4(2):bvz028.

49. Ogata T, Niihori T, Tanaka N, Kawai M, Nagashima T, Funayama R, et al. TBX1 mutation identified by exome sequencing in a Japanese family with 22q11.2 deletion syndrome-like craniofacial features and hypocalcemia. PLoS One. 2014;9(3):e91598.

50. Paylor R, Glaser B, Mupo A, Ataliotis P, Spencer C, Sobotka A, et al. Tbx1 haploinsufficiency is linked to behavioral disorders in mice and humans: implications for 22q11 deletion syndrome. Proc Natl Acad Sci U S A. 2006;103(20):7729–34.

51. Gong W, Gottlieb S, Collins J, Blescia A, Dietz H, Goldmuntz E, et al. Mutation analysis of TBX1 in non-deleted patients with features of DGS/VCFS or isolated cardiovascular defects. J Med Genet. 2001;38(12):E45.

52. Hiroi N, Zhu H, Lee M, Funke B, Arai M, Itokawa M, et al. A 200-kb region of human chromosome 22q11.2 confers antipsychotic-responsive behavioral abnormalities in mice. Proceedings of the National Academy of Sciences of the United States of America. 2005;102(52):19132–7.

53. Hiroi N, Hiramoto T, Harper KM, Suzuki G, Boku S. Mouse models of 22q11.2-associated autism spectrum disorder. Autism. 2012;S1(001):1–9.

54. Hiroi N, Takahashi T, Hishimoto A, Izumi T, Boku S, Hiramoto T. Copy Number Variation at 22q11.2: from rare variants to common mechanisms of developmental neuropsychiatric disorders. Mol Psychiatry. 2013;18:1153–65.

55. Hiroi N, Nishi A. Dimensional deconstruction and reconstruction of CNV-associated neuropsychiatric disorders. In: Pletnikov MV, Waddington JL, editors. Modeling the Psychopathological Dimensions of Schizophrenia: From Molecules to Behavior. Handbook of Behavioral Neuroscience. 23. London, UK 2015. p. 285–302.

56. Nishi A, Hiroi N. Genetic Mechanisms Emerging from Mouse Models of CNV-Associated Neuropsychiatric Disorders. In: Abel T, Nickl-Jockschat T, editors. The Neurobiology of Schizophrenia. New York: Academic Press/Elsevier; 2016. p. 397–417.

57. Suzuki G, Harper KM, Hiramoto T, Funke B, Lee M, Kang G, et al. Over-expression of a human chromosome 22q11.2 segment including TXNRD2, COMT and ARVCF developmentally affects incentive learning and working memory in mice. Human Molecular Genetics. 2009;18(20):3914–25.

58. Hiramoto T, Kang G, Suzuki G, Satoh Y, Kucherlapati R, Watanabe Y, et al. Tbx1: identification of a 22q11.2 gene as a risk factor for autism spectrum disorder in a mouse model. Hum Mol Genet. 2011;20(24):4775–85.

59. Takahashi T, Okabe S, Broin PO, Nishi A, Ye K, Beckert MV, et al. Structure and function of neonatal social communication in a genetic mouse model of autism. Mol Psychiatry. 2016;21(9):1208–14.

60. Kato R, Machida A, Nomoto K, Kang G, Hiramoto T, Tanigaki K, et al. Maternal approach behaviors toward neonatal calls are impaired by mother’s experiences of raising pups with a risk gene variant for autism. Dev Psychobiol. 2021;63(1):108–13.

61. Hiramoto T, Sumiyoshi A, Kato R, Yamauchi T, Takano T, Kang G, et al. Highly demarcated structural alterations in the brain and impaired social incentive learning in Tbx1 heterozygous mice. Molecular Psychiatry. 2025;30(5):1876–86.

62. Hiramoto T, Sumiyoshi A, Yamauchi T, Tanigaki K, Shi Q, Kang G, et al. Tbx1, a gene encoded in 22q11.2 copy number variant, is a link between alterations in fimbria myelination and cognitive speed in mice. Mol Psychiatry. 2022;27(2):929–38.

63. Nieder A, Mooney R. The neurobiology of innate, volitional and learned vocalizations in mammals and birds. Philos Trans R Soc Lond B Biol Sci. 2020;375(1789):20190054.

64. Boku S, Izumi T, Abe S, Takahashi T, Nishi A, Nomaru H, et al. Copy number elevation of 22q11.2 genes arrests the developmental maturation of working memory capacity and adult neurogenesis. Molecular Psychiatry. 2018;23(4):985–92.

65. Cao J, Spielmann M, Qiu X, Huang X, Ibrahim DM, Hill AJ, et al. The single-cell transcriptional landscape of mammalian organogenesis. Nature. 2019;566(7745):496–502.

66. Wang R, Zhang P, Wang J, Ma L, E W, Suo S, et al. Construction of a cross-species cell landscape at single-cell level. Nucleic Acids Res. 2023;51(2):501–16.

67. Marques S, van Bruggen D, Vanichkina DP, Floriddia EM, Munguba H, Varemo L, et al. Transcriptional Convergence of Oligodendrocyte Lineage Progenitors during Development. Dev Cell. 2018;46(4):504–17 e7.

68. Kessaris N, Fogarty M, Iannarelli P, Grist M, Wegner M, Richardson WD. Competing waves of oligodendrocytes in the forebrain and postnatal elimination of an embryonic lineage. Nat Neurosci. 2006;9(2):173–9.

69. Menn B, Garcia-Verdugo JM, Yaschine C, Gonzalez-Perez O, Rowitch D, Alvarez-Buylla A. Origin of oligodendrocytes in the subventricular zone of the adult brain. J Neurosci. 2006;26(30):7907–18.

70. Rushing G, Ihrie RA. Neural stem cell heterogeneity through time and space in the ventricular-subventricular zone. Front Biol (Beijing). 2016;11(4):261–84.

71. Fuentealba LC, Rompani SB, Parraguez JI, Obernier K, Romero R, Cepko CL, et al. Embryonic Origin of Postnatal Neural Stem Cells. Cell. 2015;161(7):1644–55.

72. Hiramoto T, Boku S, Kang G, Abe S, Nagashima M, Barbachan e Silva M, et al. Transcriptional regulation of neonatal neural stem cells is a determinant of social behavior. BioRxiv. 2023;11(12):468452.

73. Nishiyama A, Lin XH, Giese N, Heldin CH, Stallcup WB. Co-localization of NG2 proteoglycan and PDGF alpha-receptor on O2A progenitor cells in the developing rat brain. J Neurosci Res. 1996;43(3):299–314.

74. Rivers LE, Young KM, Rizzi M, Jamen F, Psachoulia K, Wade A, et al. PDGFRA/NG2 glia generate myelinating oligodendrocytes and piriform projection neurons in adult mice. Nat Neurosci. 2008;11(12):1392–401.

75. Sil S, Periyasamy P, Thangaraj A, Chivero ET, Buch S. PDGF/PDGFR axis in the neural systems. Mol Aspects Med. 2018;62:63–74.

76. Brandebura AN, Morehead M, Heller DT, Holcomb P, Kolson DR, Jones G, et al. Glial Cell Expansion Coincides with Neural Circuit Formation in the Developing Auditory Brainstem. Dev Neurobiol. 2018;78(11):1097–116.

77. Beiter RM, Rivet-Noor C, Merchak AR, Bai R, Johanson DM, Slogar E, et al. Evidence for oligodendrocyte progenitor cell heterogeneity in the adult mouse brain. Sci Rep. 2022;12(1):12921.

78. Arnold JS, Werling U, Braunstein EM, Liao J, Nowotschin S, Edelmann W, et al. Inactivation of Tbx1 in the pharyngeal endoderm results in 22q11DS malformations. Development. 2006;133(5):977–87.

79. Matsuo N, Takao K, Nakanishi K, Yamasaki N, Tanda K, Miyakawa T. Behavioral profiles of three C57BL/6 substrains. Front Behav Neurosci. 2010;4:29.

80. Hiroi N. Critical Reappraisal of Mechanistic Links of Copy Number Variants to Dimensional Constructs of Neuropsychiatric Disorders in Mouse Models. Psychiatry and Clinical Neurosciences. 2018;72(5):301–21.

81. Yang D, Wang Y, Qi T, Zhang X, Shen L, Ma J, et al. Phosphorylation of pyruvate dehydrogenase inversely associates with neuronal activity. Neuron. 2024;112(6):959–71 e8.

82. Readhead C, Popko B, Takahashi N, Shine HD, Saavedra RA, Sidman RL, et al. Expression of a myelin basic protein gene in transgenic shiverer mice: correction of the dysmyelinating phenotype. Cell. 1987;48(4):703–12.

83. Meschkat M, Steyer AM, Weil MT, Kusch K, Jahn O, Piepkorn L, et al. White matter integrity in mice requires continuous myelin synthesis at the inner tongue. Nat Commun. 2022;13(1):1163.

84. Arellano JI, Duque A, Rakic P. A coming-of-age story: adult neurogenesis or adolescent neurogenesis in rodents? Front Neurosci. 2024;18:1383728.

85. Bronson FH, Dagg CP, Snell GD. Reproduction. In: Green EL, editor. Biology of the Laboratory Mouse. Second Edition ed. New York: Dover Publications, Inc.; 2007. p. Oneline publication.

86. Nakamura M, Ye K, MB ES, Yamauchi T, Hoeppner DJ, Fayyazuddin A, et al. Computational identification of variables in neonatal vocalizations predictive for postpubertal social behaviors in a mouse model of 16p11.2 deletion. Mol Psychiatry. 2021;26(11):6578–88.

87. Gao Z, Daquinag AC, Su F, Snyder B, Kolonin MG. PDGFRalpha/PDGFRbeta signaling balance modulates progenitor cell differentiation into white and beige adipocytes. Development. 2018;145(1).

88. Marcelin G, Ferreira A, Liu Y, Atlan M, Aron-Wisnewsky J, Pelloux V, et al. A PDGFRalpha-Mediated Switch toward CD9(high) Adipocyte Progenitors Controls Obesity-Induced Adipose Tissue Fibrosis. Cell Metab. 2017;25(3):673–85.

89. Harper KM, Hiramoto T, Tanigaki K, Kang G, Suzuki G, Trimble W, et al. Alterations of social interaction through genetic and environmental manipulation of the 22q11.2 gene Sept5 in the mouse brain. Human Molecular Genetics. 2012;21(15):3489–99.

90. Suzuki G, Harper KM, Hiramoto T, Sawamura T, Lee M, Kang G, et al. Sept5 deficiency exerts pleiotropic influence on affective behaviors and cognitive functions in mice. Human Molecular Genetics. 2009;18(9):1652–60.

91. Yamauchi T, Kang G, Hiroi N. Heterozygosity of murine Crkl does not recapitulate behavioral dimensions of human 22q11.2 hemizygosity. Genes Brain and Behavior. 2020;20(5):e12719.

92. Paylor R, Spencer CM, Yuva-Paylor LA, Pieke-Dahl S. The use of behavioral test batteries, II: effect of test interval. Physiol Behav. 2006;87(1):95–102.

93. Fonseca AH, Santana GM, Bosque Ortiz GM, Bampi S, Dietrich MO. Analysis of ultrasonic vocalizations from mice using computer vision and machine learning. Elife. 2021;10.

94. Sakamoto Y, Takano T, Shimoyama S, Hiramoto T, Hiroi N, Nakamura K. Prepartum bumetanide treatment reverses altered neonatal social communication but non-specifically reduces post-pubertal social behavior in a mouse model of fragile X syndrome. Genomic Psychiatry. 2025;1(1):61–72.

95. Kaiser T, Allen HM, Kwon O, Barak B, Wang J, He Z, et al. MyelTracer: A Semi-Automated Software for Myelin g-Ratio Quantification. eNeuro. 2021;8(4).

96. Cioffi S, Martucciello S, Fulcoli FG, Bilio M, Ferrentino R, Nusco E, et al. Tbx1 regulates brain vascularization. Hum Mol Genet. 2014;23(1):78–89.

97. Cioffi S, Flore G, Martucciello S, Bilio M, Turturo MG, Illingworth E. VEGFR3 modulates brain microvessel branching in a mouse model of 22q11.2 deletion syndrome. Life Sci Alliance. 2022;5(12).

98. Allen Brain Atlas. Mouse Brain. http://wwwbrain-maporg. 2017.

99. Nait Oumesmar B, Vignais L, Baron-Van Evercooren A. Developmental expression of platelet-derived growth factor alpha-receptor in neurons and glial cells of the mouse CNS. J Neurosci. 1997;17(1):125–39.

100. Zhu X, Bergles DE, Nishiyama A. NG2 cells generate both oligodendrocytes and gray matter astrocytes. Development. 2008;135(1):145–57.

101. Marques S, Zeisel A, Codeluppi S, van Bruggen D, Mendanha Falcao A, Xiao L, et al. Oligodendrocyte heterogeneity in the mouse juvenile and adult central nervous system. Science. 2016;352(6291):1326–9.

102. Ozarkar SS, Patel RKR, Vulli T, Friar CA, Burette AC, Philpot BD. Regional analysis of myelin basic protein across postnatal brain development of C57BL/6J mice. Front Neuroanat. 2025;19:1535745.

103. Bastedo WE, Scott RW, Arostegui M, Underhill TM. Single-cell analysis of mesenchymal cells in permeable neural vasculature reveals novel diverse subpopulations of fibroblasts. Fluids Barriers CNS. 2024;21(1):31.

104. Takeuchi A, Takahashi Y, Iida K, Hosokawa M, Irie K, Ito M, et al. Identification of Qk as a Glial Precursor Cell Marker that Governs the Fate Specification of Neural Stem Cells to a Glial Cell Lineage. Stem Cell Reports. 2020;15(4):883–97.

105. Dani N, Herbst RH, McCabe C, Green GS, Kaiser K, Head JP, et al. A cellular and spatial map of the choroid plexus across brain ventricles and ages. Cell. 2021;184(11):3056–74 e21.

106. Downes N, Mullins P. The development of myelin in the brain of the juvenile rat. Toxicol Pathol. 2014;42(5):913–22.

107. Thompson CL, Ng L, Menon V, Martinez S, Lee CK, Glattfelder K, et al. A high-resolution spatiotemporal atlas of gene expression of the developing mouse brain. Neuron. 2014;83(2):309–23.

108. Redmond SA, Figueres-Onate M, Obernier K, Nascimento MA, Parraguez JI, Lopez-Mascaraque L, et al. Development of Ependymal and Postnatal Neural Stem Cells and Their Origin from a Common Embryonic Progenitor. Cell Rep. 2019;27(2):429–41 e3.

109. O’Rourke M, Cullen CL, Auderset L, Pitman KA, Achatz D, Gasperini R, et al. Evaluating Tissue-Specific Recombination in a Pdgfralpha-CreERT2 Transgenic Mouse Line. PLoS One. 2016;11(9):e0162858.

110. Kang SH, Fukaya M, Yang JK, Rothstein JD, Bergles DE. NG2+ CNS glial progenitors remain committed to the oligodendrocyte lineage in postnatal life and following neurodegeneration. Neuron. 2010;68(4):668–81.

111. Tripathi RB, Rivers LE, Young KM, Jamen F, Richardson WD. NG2 glia generate new oligodendrocytes but few astrocytes in a murine experimental autoimmune encephalomyelitis model of demyelinating disease. J Neurosci. 2010;30(48):16383–90.

112. Komitova M, Zhu X, Serwanski DR, Nishiyama A. NG2 cells are distinct from neurogenic cells in the postnatal mouse subventricular zone. J Comp Neurol. 2009;512(5):702–16.

113. Rosenberg AB, Roco CM, Muscat RA, Kuchina A, Sample P, Yao Z, et al. Single-cell profiling of the developing mouse brain and spinal cord with split-pool barcoding. Science. 2018;360(6385):176–82.

114. Yao Z, Liu H, Xie F, Fischer S, Adkins RS, Aldridge AI, et al. A transcriptomic and epigenomic cell atlas of the mouse primary motor cortex. Nature. 2021;598(7879):103–10.

115. Thompson PM, Jahanshad N, Ching CRK, Salminen LE, Thomopoulos SI, Bright J, et al. ENIGMA and global neuroscience: A decade of large-scale studies of the brain in health and disease across more than 40 countries. Transl Psychiatry. 2020;10(1):100.

116. Kochunov P, Hong LE, Dennis EL, Morey RA, Tate DF, Wilde EA, et al. ENIGMA-DTI: Translating reproducible white matter deficits into personalized vulnerability metrics in cross-diagnostic psychiatric research. Hum Brain Mapp. 2020;43(1):194–206.

117. Koshiyama D, Fukunaga M, Okada N, Morita K, Nemoto K, Usui K, et al. White matter microstructural alterations across four major psychiatric disorders: mega-analysis study in 2937 individuals. Mol Psychiatry. 2020;25(4):883–95.

118. Kelly S, Jahanshad N, Zalesky A, Kochunov P, Agartz I, Alloza C, et al. Widespread white matter microstructural differences in schizophrenia across 4322 individuals: results from the ENIGMA Schizophrenia DTI Working Group. Mol Psychiatry. 2018;23(5):1261–9.

119. Villalon-Reina JE, Martinez K, Qu X, Ching CRK, Nir TM, Kothapalli D, et al. Altered white matter microstructure in 22q11.2 deletion syndrome: a multisite diffusion tensor imaging study. Mol Psychiatry. 2020;25(11):2818–31.

